# Probabilistic solvers enable a straight-forward exploration of numerical uncertainty in neuroscience models

**DOI:** 10.1101/2021.04.27.441605

**Authors:** Jonathan Oesterle, Nicholas Krämer, Philipp Hennig, Philipp Berens

## Abstract

Understanding neural computation on the mechanistic level requires models of neurons and neuronal networks. To analyze such models one typically has to solve coupled ordinary differential equations (ODEs), which describe the dynamics of the underlying neural system. These ODEs are solved numerically with deterministic ODE solvers that yield single solutions with either no, or only a global scalar bound on precision. It can therefore be challenging to estimate the effect of numerical uncertainty on quantities of interest, such as spike-times and the number of spikes. To overcome this problem, we propose to use recently developed sampling-based probabilistic solvers, which are able to quantify such numerical uncertainties. They neither require detailed insights into the kinetics of the models, nor are they difficult to implement. We show that numerical uncertainty can affect the outcome of typical neuroscience simulations, e.g. jittering spikes by milliseconds or even adding or removing individual spikes from simulations altogether, and demonstrate that probabilistic solvers reveal these numerical uncertainties with only moderate computational overhead.

## 1 Introduction

Computational neuroscience is built around computational models of neurons that allow the simulation and analysis of signal processing in the central nervous system. These models can describe neural computations on different levels of abstraction. On the *statistical level*, e.g. generalized linear models have been used to provide a probabilistic model mapping environmental variables to neural activity [1]. For such statistical models, quantifying the uncertainty of the parameters can be achieved using Bayesian approaches [2]. On the *mechanistic level*, the models typically take the form of systems of coupled ordinary differential equations (ODEs), which describe the dynamics of the membrane potential and give rise to the spike-times [3, 4]. Recently, likelihood-free inference approaches have made it possible to perform uncertainty-aware inference even for such complicated mechanistic models [5–7].

However, mechanistic models of neurons are subject to an additional source of uncertainty: the numerical error caused by the solution of the model’s ODEs with a concrete algorithm [8]. This arises because all numerical solvers are necessarily run with finite time and limited resources, so their estimate diverges from the true solution of the ODE, even if the problem is well-posed. When simulating neurons, one would like to compute a numerical solution close to the true solution of the ODE, to ensure that conclusions drawn from the simulations are based on the mechanisms described by the model rather than the specific choice, setting and implementation of the ODE solver.

Many of the well-established numerical solvers do report a global error estimate and a corresponding tolerance that can be set by the user [9, Chapter II.4]. This global scalar error, though, does not capture how the numerical error arising from finite step-sizes used in practice affects crucial quantities of interest in the simulation, such as spike-times or the number of spikes. In practice, it can therefore be challenging to select a tolerance that strikes a good balance between run time and accuracy.

For some of the most common mechanistic models in neuroscience like the Hodgkin-Huxley or Izhikevich neuron model, errors in numerical integration have been studied in detail for a range of solvers and different integration step-sizes [10–12]. These studies have shown that standard solvers are often not the best choice in terms of accuracy or the accuracy vs. run time tradeoff. Therefore, the authors of these studies proposed to use specific solvers for the analyzed models, e.g. the Parker-Sochacki method for the Hodgkin-Huxley and Izhikevich neuron [10], an exponential midpoint method [11] or second-order Strang splitting [12] for Hodgkin-Huxley-like models. While improving computations for the specific problems, applying these to other scenarios requires a detailed understanding of the kinetics of the neuron model of interest; and while choosing a “good” solver for a given model is important, it is typically not necessary to choose the “best” ODE solver. In many cases, it can be sufficient to ensure that the computed solution is within a certain accuracy.

As a more general approach to quantify the numerical uncertainty in mechanistic models in neuroscience, we therefore propose to use probabilistic ODE solvers [8, 13, 14]. In contrast to classical ODE solvers, this class of solvers does not only yield a single solution, but instead a distribution over solutions that quantifies the numerical uncertainty.

Several frameworks for probabilistic ODE solvers have been proposed, which differ mostly in the tradeoff between computational cost and flexibility of the posterior, from fast Gaussian filters [15–17] to sampling-based approaches [18–22]. These solvers have been mostly tested for well-behaved systems with well-behaved solutions, but the ODEs used to simulate neural activity model the non-linear membrane dynamics that underlie the all-or-none nature of an action potential. Here, we use two related approaches of probabilistic ODE integration, the state perturbation proposed by Conrad et al. [18] and the step-size perturbation of Abdulle and Garegnani [22]. Both build on existing explicit, iterative ODE solvers and stochastically perturb the numerical integration of individual steps taken by the underlying solvers. These perturbations make the solution of every step probabilistic and therefore of the solution as a whole. The magnitude of the perturbation has to be calibrated, such that the solver’s output distribution reflects the numerical uncertainty in the solution.

Here, we explore the potential of probabilistic ODE solvers for neuron models. We show how probabilistic solvers can be used to quantify and reveal numerical uncertainty caused by the numerical ODE integration and demonstrate that the solver outputs are easy to interpret. For this, we simulate typical neuron models, namely the Izhikevich neuron model [23], as a representative of leaky-integrate-and-fire neuron models, single-compartment Hodgkin-Huxley models [24] and a model with three synaptically coupled Hodgkin-Huxley-like neurons [25] as an example of a neuronal network model. Lastly, we discuss practical considerations and limitations of probabilistic solvers such as the calibration of the perturbation and the computational overhead.

Taken together, our results suggest that probabilistic ODE solvers should be considered as a useful tool for the simulation of neuronal systems, to increase the quality and reliability of such simulations over those achieved with classic solvers.

## 2 Methods and Models

### 2.1 Probabilistic solvers

Simulating neuron models typically amounts to solving an initial value problem (IVP) based on a set of coupled ODEs. In abstract form, an initial value problem is given by

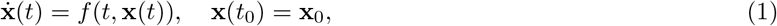

where *f*, **x**_0_ and *t*_0_ are known and **x**(*t*) for *t > t*_0_ is the quantity of interest. The solution to the initial value problem at time *t* + Δ*t* provided the solution at time *t*, is given by integrating Eq. (1) from *t* to *t* + Δ*t*:

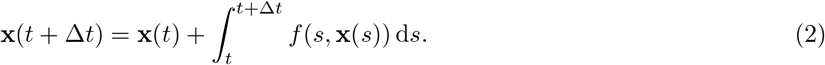

Except for special cases, this integral has no analytic form and must be solved numerically. For example, the forward Euler method approximates the integral as 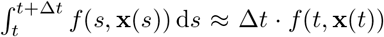. To simulate a neuron, Eq. (2) is solved iteratively, which results in a sequence of solutions *X* = [*x*(*t*_0_), *x*(*t*_1_), *x*(*t*_2_), *…, x*(*t*_*N*_)] for a set of time points with *t*_*i*+1_ *> t*_*i*_ and a maximum time point *t*_*N*_. Standard solvers yield a deterministic solution in every step, and therefore for the solution *X* as a whole. In contrast, the probabilistic solvers used in this study stochastically perturb the numerical integration used to approximate Eq. (2), which makes the solution of every step—and therefore of the whole solution—probabilistic. For a given IVP and solver, one can therefore generate a sample distribution of solutions *X* by repeating the iterative numerical integration from *t*_0_ to *t*_*N*_ multiple times. To create these probabilistic solvers, we implemented the state perturbation algorithm of Conrad et al. [18] and the step-size perturbation algorithm of Abdulle and Garegnani [22].

In the state perturbation algorithm [18], in each step of the numerical integration, a small i.i.d. noise term ***ξ***_*t*_ is added to the solution **x**_det_(*t* + Δ*t*) of a corresponding deterministic integration scheme:

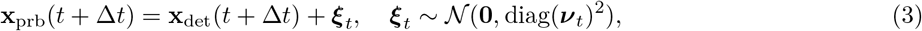

where ***ν***_*t*_ controls the magnitude of the perturbation. The perturbation is only efficient when ***ν***_*t*_ is of the right order: if chosen too small, the uncertainty will be underestimated; if chosen too large, it will render the solver output useless. Conrad et al. [18] suggested calibrating ***ν***_*t*_ to replicate the amount of error introduced by the numerical scheme. We chose ***ν***_*t*_ = *σ****ε***_*t*_ using the error estimator ***ε***_*t*_ readily available in methods that were developed for step-size adaptation (see Appendix A), and a scalar perturbation parameter *σ* that can be adjusted to calibrate the perturbation. If not stated otherwise, we used *σ*=1. An example of this perturbation method is shown in Fig. 1A for a single integration step and in Fig. 1C for an Izhikevich neuron model.

**Figure 1:**
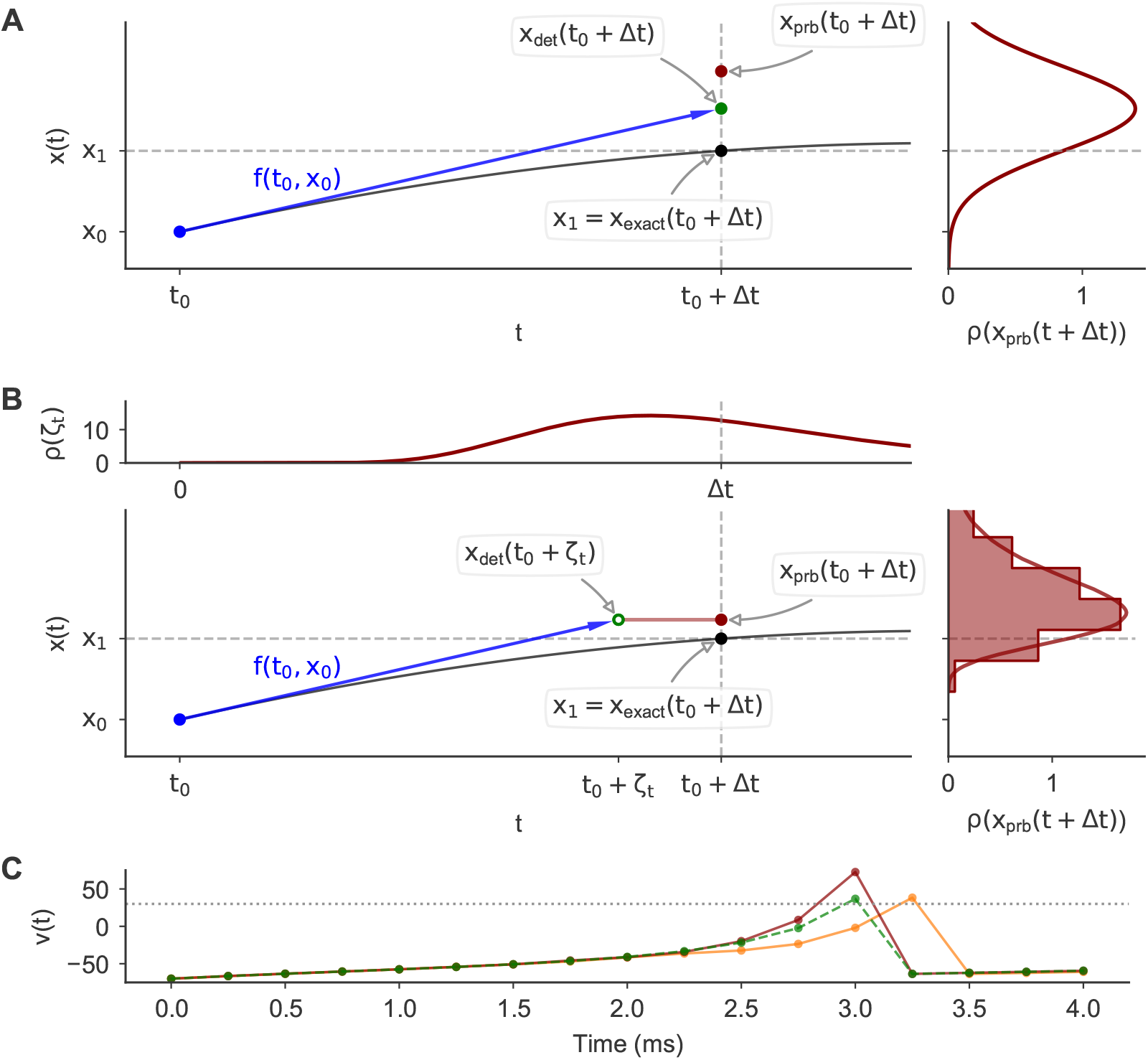
Illustration of probabilistic ODE solvers. **(A)** *Left* : A single integration step with a probabilistic forward Euler method using the state perturbation method [18] for the ODE *f* (*t, x*(*t*)) = 3 *· x*(*t*) *·* sin(*t* + 3) and the exact solution *x*(*t*) = exp(−3 *·* cos(*t* + 3)) (black curve). We set *t*_0_ = 0 and the step-size to Δ*t*=0.1. The exact solution at *t* = *t*_0_ + Δ*t* is highlighted (black dot). A first order solution is computed using forward Euler: *x*_det_(*t* + Δ*t*) = *x*_0_ + Δ*t · f* (*t*_0_, *x*_0_) (*x*_det_: green dot, *f* : blue arrow). *Right* : The probability density function *ρ* of *x*_prb_(*t* + Δ*t*), where *x*_prb_(*t* + Δ*t*) is the output of the probabilistic step. In the state perturbation, *ρ* is a normal distribution with mean *x*_det_(*t* + Δ*t*) and a standard deviation based on a local error estimator (see Section 2.1). A random sample is shown for illustration (red dot). **(B)** Similar to (A), but for the step-size perturbation method [22] using a log-normal perturbation distribution. Instead of integrating from *t*_0_ to *t*_0_ + Δ*t*, the ODE is integrated from *t*_0_ to *t*_0_ + *ζ*_*t*_, where *ζ*_*t*_ is randomly drawn from a log-normal distribution (top panel). The solution of this perturbed integration *x*_det_(*t* + *ζ*_*t*_) (green circle) is then used as the solution *x*_prb_(*t* +Δ*t*) of the probabilistic step (red dot), making *x*_prb_(*t* + Δ*t*) a random variable with a distribution *ρ*(*x*_prb_(*t* + Δ*t*)) (right panel), that has no general analytical form but is dependent on the ODE and the solver. Therefore, *ρ*(*x*_prb_(*t* +Δ*t*)) is shown as an empirical histogram and a kernel density estimate. **(C)** Simulations of an Izhikevich neuron with a deterministic (green dashed line) and a probabilistic forward Euler method using state perturbation (two samples: red and orange).

A related approach to stochastically perturbing the numerical integration was proposed by Abdulle and Garegnani [22], where noise is added to the integration step-size (i.e. to the “input” of the solver, rather than the “output”, cf. Fig. 1B). The numerical integration is performed using the perturbed step-size *ζ*_*t*_, but the computed solution is treated as the solution for the original step-size Δ*t*:

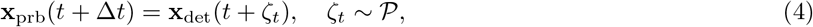

where *ζ*_*t*_ is the i.i.d. perturbed step-size drawn from a distribution 𝒫 and **x**_det_(•) is a deterministic integration scheme that approximates Eq. (2). For example, for the forward Euler method Eq. (4) would be computed as **x**_prb_(*t* + Δ*t*) = **x**_det_(*t*) + *ζ*_*t*_ *· f* (*t*, **x**_det_(*t*)). Abdulle and Garegnani [22] defined three properties the i.i.d. random variables *ζ*_*t*_ should fulfill:

- *P* (*ζ*_*t*_ *>* 0) = 1, where *P* is the probability,
- there exists Δ*t* such that 𝔼[*ζ*_*t*_] = Δ*t*, and
- there exist *p* ≥ 0.5 and *C >* 0 independent of *t* such that 𝔼[(*ζ*_*t*_ − Δ*t*)^2^] = *C ·* Δ*t*^2*p*+1^. Based on these restrictions, they proposed, as an example, to use a log-normal distribution:

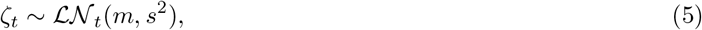

where the mean *m* and standard deviation *s* of the underlying normal distribution should be chosen such that 𝔼[*ζ*_*t*_] = Δ*t* and 𝔼[(*ζ*_*t*_ − Δ*t*)^2^] = *C ·* Δ*t*^2*p*+1^ hold for some *C >* 0 and *p* ≥ 0.5 independent of Δ*t. m* and *s* can therefore be defined as:

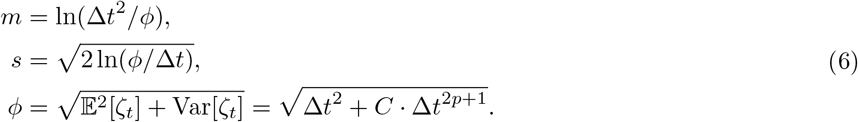

Using *p* = *O*, where *O* is the order of the method, ensures that the mean-squared convergence order of the method is not changed. We therefore used *p* = *O* throughout. We further generalized the example provided by Abdulle and Garegnani [22] in which *C* = 1 to a parametrized distribution by setting *C* = *σ*^2^, i.e. setting 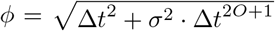. The introduction of the perturbation parameter *σ* allows to—similarly to the perturbation parameter used in the state-perturbation—adjust and calibrate the magnitude of perturbation. If not stated otherwise, we used *σ*=1. The perturbation is illustrated in Fig. 1B for the first order forward Euler scheme.

### 2.2 Choice of solvers

We used the perturbation methods described above to create probabilistic versions of the solvers listed in Table 1.

**Table 1:** Summary of the ODE solvers used in this paper.

| Abbr. | $O$ | $O_e$ | Method & Error estimate |
| --- | --- | --- | --- |
| FE | 1 | 2 | *Forward Euler* with Heun’s method for error estimation. |
| EE | 1 |  | *Exponential Euler*. |
| EEMP | 2 |  | *Exponential Euler Midpoint* [11]. |
| RKBS | 3 | 2 | *Bogacki–Shampine*, an embedded Runge-Kutta method [26]. |
| RKCK | 4 | 5 | *Cash–Karp method*, an embedded Runge-Kutta method [27]. |
| RKDP | 5 | 4 | *Dormand–Prince*, an embedded Runge-Kutta method [28]. |

$O$ and $O_e$ are the orders of the solution and the error estimator, respectively. See Appendix B for details.

The usage of fixed (f) and adaptive (a) step-sizes is indicated with subscripts, and the perturbation method is indicated using the superscripts—*x* for the state perturbation [18] and *t* for the step-size perturbation [22]— meaning that e.g. 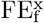 is referring to a forward Euler method using fixed step-sizes and the state perturbation. For the exponential integrators, we chose to only use the step-size perturbation because it preserves the important property of these solvers that the activation and inactivation variables can not leave the interval [0, 1], and also because there are no established methods for local error estimation for these methods.

The second order exponential integrator EEMP was implemented based on the version by Börgers and Nectow [11] (Appendix B), which is a modification of the midpoint method by Oh and French [29]. Computation of Runge-Kutta steps and step-size adaptation were based on the respective scipy implementations [30]. To avoid computational overhead, we only computed the local error estimates when necessary, i.e. for adaptive step-sizes or the state perturbation.

### 2.3 Interpolation

The iterative solvers used in this study yield solutions for **x**(*t*) on either a fixed and equidistant grid of time points *T* or, in the case of adaptive step-size solvers, on a finite set of time points *T* automatically chosen by the solver. To interpolate these solutions for example for spike-time estimation (see Section 2.4), we used linear interpolation for FE, EE and EEMP between solutions of single steps. To interpolate the steps of the Runge-Kutta methods we utilized the “dense output” implemented in the respective scipy methods [30]. These “dense outputs” allow to evaluate the solution between two steps **x**(*t*_*i*_) and **x**(*t*_*i*+1_) for any *t* with *t*_*i*_ ≤ *t* ≤ *t*_*i*+1_ without any additional ODE evaluation. To not discard the effect of the state perturbation during interpolation, we defined the dense output 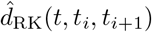 for a state perturbed Runge-Kutta step from time *t*_*i*_ to *t*_*i*+1_ as:

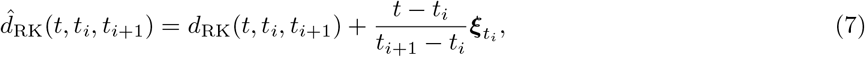

where *d*_RK_(*t, t*_*i*_, *t*_*i*+1_) is the dense output of the respective deterministic Runge-Kutta step and 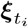 is the perturbation noise that was added to this step to compute **x**(*t*_*i*+1_) (see Eq. (3). This is a simplified version of the continuous-time output proposed by Conrad et al. [18].

### 2.4 Spike-time estimation

To determine spike-times based on simulated voltage traces *v*(*t*), we interpolated the ODE solutions for all steps where *v*(*t*) started from below and ended above a certain threshold voltage *v*_th_. For lineally interpolated solutions (Section 2.3) we computed spike-times as follows. For every step from a time *t*_*i*_ to *t*_*i*+1_ with *v*(*t*_*i*_) *< v*_th_ ≤ *v*(*t*_*i*+1_) we estim ated the respective spike-time *t*_spike_ as:

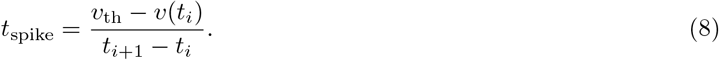

To estimate spike-times for Runge-Kutta methods with “dense-outputs”, we utilized scipy’s “brentq” root finding algorithm to determine the time point *t*_spike_ when the threshold is reached, i.e. |*v*(*t*_spike_)−*v*_th_| *< ε* with *ε* = 1*e*−12.

### 2.5 Common ODE models in computational neuroscience

In this study, we use probabilistic ODE solvers to analyze the effect of numerical uncertainty in the following neuroscience models:

- The Izhikevich neuron model with a wide range of dynamics,
- the Hodgkin-Huxley neuron model,
- and a small network of Hodgkin-Huxley neurons.

We picked these models to cover both single neuron models and models of neuronal networks.

#### 2.5.1 Single Izhikevich neurons

The Izhikevich neuron (IN) model is a simplified non-linear single neuron model that has been used e.g. to build large-scale models of the brain [31] and to understand oscillatory phenomena in the cortex [32,33] and the olfactory bulb [34]. An attractive property of the IN is that a whole range of different response dynamics can be simulated (Fig. S1) depending on the setting of the parameters ***θ*** = [*a, b, c, d*] [23]. The IN is described by the following pair of ODEs [32]:

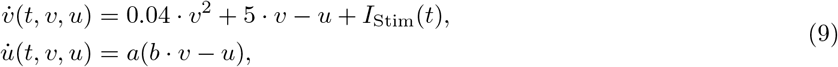

where *v* is the membrane potential, *u* is a recovery variable and *I*_Stim_ is a given input current. Whenever the threshold is reached, i.e. *v*(*t*) ≥ 30, a “spike” is triggered and the neuron is reset in the next time step of the simulation:

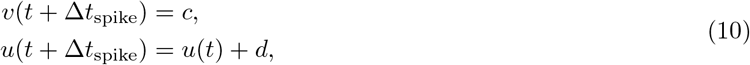

where Δ*t*_spike_ ≥ 0. Typically Δ*t*_spike_ = Δ*t* is used, but to facilitate the comparison between different stepsizes we used Δ*t*_spike_ = 0 instead. The reset is problematic, because it introduces an error of order *O*(Δ*t*) [10], independent of the solver scheme. This is because spikes can only occur after a full step of integration and the value *u*(*t* + Δ*t*_spike_) in Eq. (10) is dependent on the previous value *u*(*t*).

To address this problem, we implemented two complementary strategies. Fist, we adapted Eq. (9) such that whenever 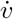 and 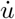 would have been evaluated for *v*(*t*) ≥ 30—which can only happen for multi-stage methods—the derivatives were evaluated for *v*(*t*) = 30 instead. Second, we implemented the strategy suggested by Stewart et al. [10]: Every step resulting in a reset is split into two intermediate steps, a step until the threshold is reached, and a step after the reset. For this, the spike-time *t*_spike_ during such as step was estimated as described in Section 2.4 with a threshold of *v*_th_ = 30. Then, the pre-reset step solution **x**(*t*_spike_) was approximated based on the interpolation strategies described in Section 2.3. And finally, the post-reset step solution 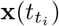 was computed by resetting (see Eq. (10)) and integrating **x** from *t*_spike_ 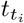.

#### 2.5.2 Single Hodgkin-Huxley neurons

Hodgkin-Huxley (HH) models [24] are widely used to simulate single and multi-compartment neurons. We study both the classical HH neuron [24] and a single compartment HH-like neuron model [35] prominently used to study the stomatogastric ganglion (STG) [25]. Both models are described by ODEs including the membrane potential *v*(*t*) described by:

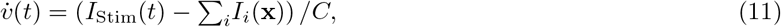

where *C* is the membrane capacitance, *I*_Stim_ is the stimulation current and *I*_*i*_ are membrane currents. These membrane currents are described by the following equation:

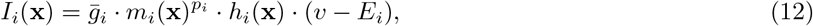

where *E*_*i*_ is the reversal potential of the current, 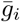 is the maximum channel conductance, *p*_*i*_ are integer exponents, and *m*_*i*_ and *h*_*i*_ are activation and inactivation functions. *m*_*i*_ and *h*_*i*_ were modeled by the following differential equations:

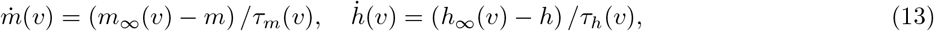

where *m*_∞_, *τ*_*m*_, *h*_∞_, and *τ*_*h*_ are voltage dependent functions defining the channel’s kinetics. For non-inactivating channels, *h*_*i*_ is removed from Eq. (12). In the classical HH model, this amounts to a 4-dimensional ODE [36]. For the STG neuron, which has eight instead of two membrane currents and also implements a model for the intracellular calcium concentration, the ODE is 13-dimensional [35]. The respective parametrizations can be found in Appendix C.

We simulated the HH neuron’s response to two different input current *I*_Stim_, a step and a noisy step stimulus. Both stimuli were 200 ms long, with *I*_Stim_(*t*) = 0 for *t < t*_onset_ and *t* ≥ *t*_offset_, where *t*_onset_ = 10 ms and *t*_offset_ = 190 ms. The amplitude of the step stimulus for *t*_onset_ ≥ *t < t*_offset_ was *I*_Stim_(*t*) = 0.2 mA. The amplitude of the noisy step stimulus were created by drawing 99 values from a uniform distribution between 0.0 mA and 0.4 mA that were spaced equidistantly between *t*_onset_ and *t*_offset_. These points were interpolated using a cubic spline with endpoints at *t*_onset_ and *t*_offset_. At the endpoints both *I*_Stim_ and its derivative were set to zero. The single STG neuron was simulated for 3 s using a step stimulus starting at *t*_onset_ = 0.9 s with an amplitude of *I*_Stim_(*t*) = 3 nA.

#### 2.5.3 STG model

The STG neuron model described above was used by Prinz et al. [25] in a network of three synaptically coupled neurons, ABPD, LP and PY, to study their firing patterns in dependence of the synaptic and neuronal parametrizations. In the model, there are seven synapses connecting the neurons, that are either modeled as slow or fast synapses. The postsynaptic input current *I*_*i*_ to a neuron is described by:

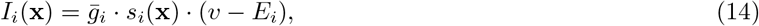

where, similarly to Eq. (12), *E*_*i*_ is the reversal potential of the current, 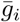 is the synapse’s maximum conductance, *v* is the membrane potential of the postsynaptic neuron and *s* is the activation function of the synapse. *s* is described by the following differential equation:

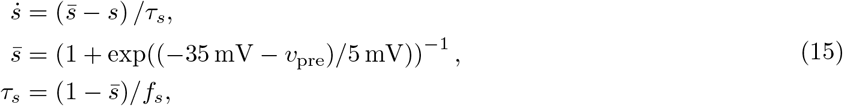

where *v*_pre_ is the membrane potential of the presynaptic neuron and *τ*_*s*_ and *f*_*s*_ are constants (see Appendix C).

### 2.6 Quantifying numerical uncertainty

#### 2.6.1 Reference solutions

None of the aforementioned neuron models has an analytical solution. It is therefore not possible to compare simulations to the true solutions of the respective IVPs. As a substitute, we computed reference solutions using a deterministic RKDP_a_ solver with a tolerance of *κ*=1e − 12 and a maximum step-size dependent on the model investigated (0.01 ms for IN and HH; 0.1 ms for the STG model). To obtain a reference solution at the same time points of a given fixed step-size solution *X* = [**x**(*t*_0_), …, **x**(*t*_*M*_)], we forced the reference solver to evaluate **x**(*t*) at least at all time points *T* = [*t*_0_, …, *t*_*M*_] of the given solution. For this, in every step in which the adaptive reference solver automatically picked a step-size that would skip any *t*_*i*_ in *T* by taking a too large step-size Δ*t*_*i*−1_, the step-size Δ*t*_*i*−1_ was clipped such that the step was evaluated exactly at **x**(*t*_*i*_). All solutions **x**(*t*) for *t* not in *T* were dropped before the comparison. To compare adaptive step-size solvers to reference solutions, we also forced these solvers to evaluate time points on a grid *T* = [*t*_0_, …, *t*_*M*_] with time points space equidistantly using a distance of 1 ms.

#### 2.6.2 Distance metrics

To estimate the uncertainty for a given neuron model and solver, we computed multiple solutions (samples) with the same probabilistic solver to obtain a distribution of solutions. Based on these sample distributions and the respective reference solutions, we evaluated the distributions of sample-sample distances and sample-reference distances. For this, we computed the Mean Absolute Error (MAE) between single traces of the ODE solutions as a distance measure. If not stated otherwise, MAEs were computed on the simulated membrane potentials, because this is typically the quantity of interest. For two traces of equal size **a** = [*a*_0_, …, *a*_*M*_] and **b** = [*b*_0_, …, *b*_*M*_] the MAE was defined as:

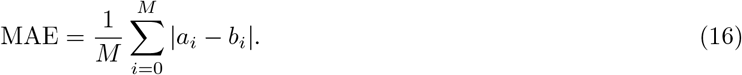

For *n* samples from a probabilistic solver, we computed the sample-sample distance distribution MAE_SM_ as the *n* MAEs between single samples and the mean trace of the other *n* − 1 samples. Sample-reference distance distributions MAE_SR_ were computed as the *n* MAEs between single samples and the reference solution. In some cases, we also computed the distance between the solution of a corresponding deterministic solver to the reference solution, abbreviated as MAE_DR_.

### 2.7 Code and availability

The probabilistic solvers and models were implemented in Python and Cython. The code is available at https://github.com/berenslab/neuroprobnum.

## 3 Results

In this study, we explored the potential of probabilistic ODE solvers in computational neuroscience. First, we study the effect of numerical uncertainty on simulations of neuron models and qualitatively show that probabilistic solvers can reveal this uncertainty in a way that is easy to interpret. Second, we provide examples and guidelines were probabilistic solvers can be useful when conducting a new study. Third, we analyze potential drawbacks of probabilistic solvers, such as computational overhead.

### 3.1 Probabilistic solvers can reveal numerical uncertainty in neuron models

To demonstrate the effect of numerical uncertainty on simulations of single neuron models, we first simulated the classical HH neuron with the step stimulus (Fig. 2A). We computed solutions with a deterministic and probabilistic EE solver for a step-size of Δ*t*=0.25 ms. Additionally, we computed a reference solution. We found that the exact spike-times of the deterministic EE solver differed substantially from the reference solution (spike-time difference 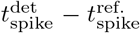 of the first three spikes: 0.7 ms, 2.8 ms, 4.5 ms). The probabilistic solver revealed this numerical uncertainty with spike-times varying substantially between samples (standard deviation (SD) of the spike-time *t*_spike_ for the first three spikes over all 20 samples: 0.2 ms, 0.9 ms, 1.2 ms).

**Figure 2:**
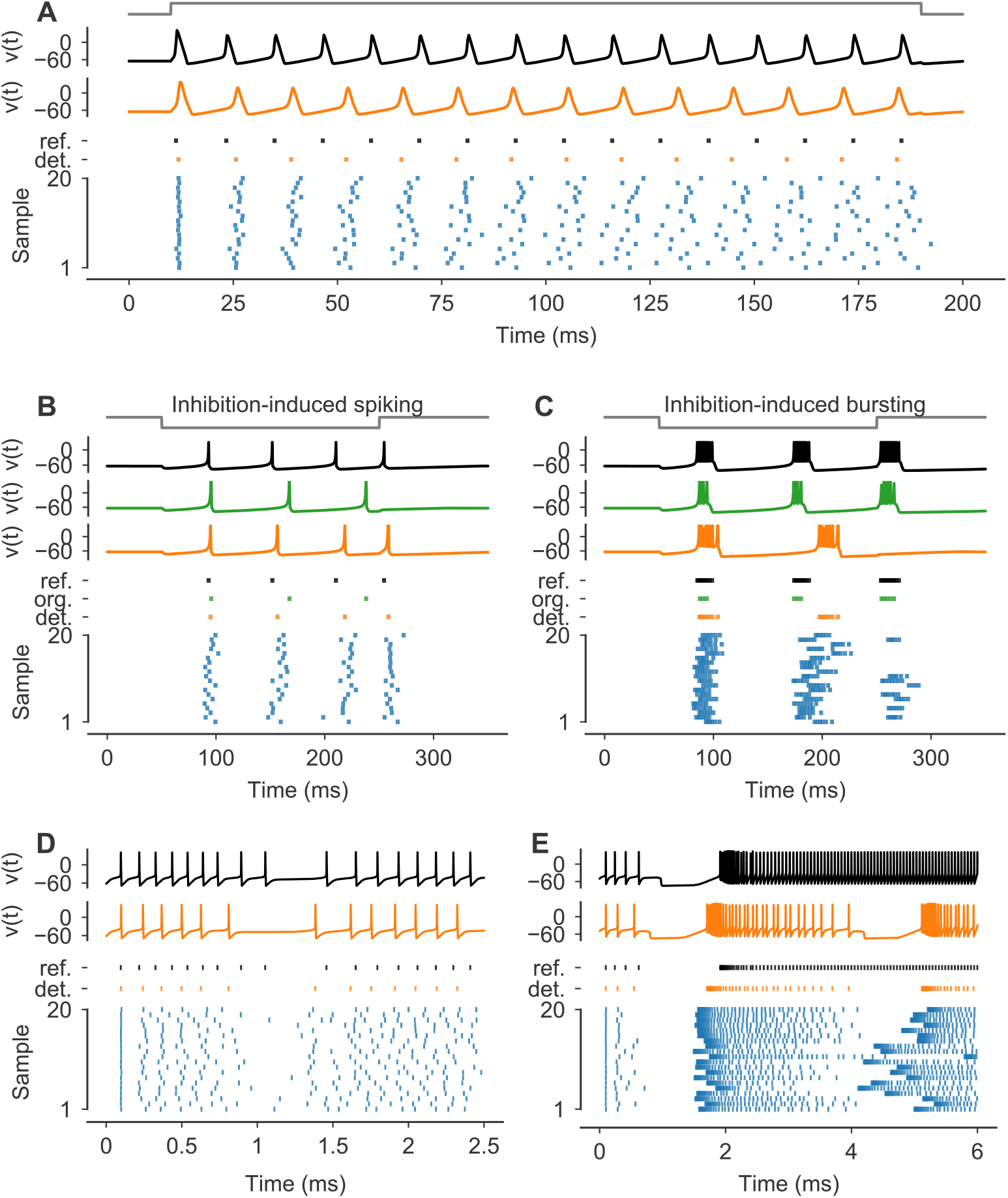
Neuron simulations can be subject to substantial numerical uncertainty. **(A)** Simulations of the classical HH model for the step stimulus *I*_Stim_ (normalized stimulus in gray). Solutions for *v*(*t*) are shown for a reference solver (black) and a deterministic EE solver with Δ*t*=0.25 ms (orange). *Bottom panel* : Spike-times of the reference (black), the deterministic EE solution (orange) and for 20 samples from a probabilistic 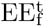 solver with Δ*t*=0.25 ms (blue). **(B, C)** Simulations of the IN model for two different parametrizations *θ* (see Appendix C) and stimuli *I*_Stim_ (normalized stimuli in gray). Solutions for *v*(*t*) are shown for a reference solver (black), the original solver scheme (green) and a deterministic FE solver (orange). Based on the original publication, the step-size Δ*t* was set to 0.5 ms for all but the reference solver. For plotting, *v*(*t*) were clipped at 30. *Bottom panels*: Spike-times are shown for the reference (black), the original solver solution (green), the deterministic FE solution (orange) and for 20 samples from a probabilistic 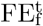 solver (blue). Samples were sorted by the number of spikes. **(D, E)** Simulations of the STG model for the two different synaptic parametrizations (see Appendix C) 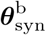 and 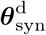, respectively. Solutions for the membrane potential *v*(*t*) of the LP neuron are shown for a reference solver (black) and a deterministic EE solver with Δ*t*=0.1 ms (orange). *Bottom panels*: Spike-times of the LP neuron are shown for the reference (black), the deterministic EE solution (orange) and for 20 samples from a probabilistic 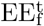 solver with Δ*t*=0.1 ms (blue). Samples were sorted by the number of spikes.

Next we simulated single INs with different parametrizations *θ*_*i*_ and response dynamics [23]. Using the original step-sizes Δ*t* and input currents *I*_*i*_, we compute solutions with the original solver scheme—which is related to a FE_f_ solver (Appendix B)—a deterministic FE scheme and a probabilistic 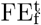 solver. We found, that for the “Inhibition-induced spiking” neuron all solvers produced similar spiking patterns in response to a negative current step (Fig. 2B). However, the original solver produced longer intervals between the spikes compared to the reference, resulting in only three instead of four spikes. The deterministic FE solution matched the reference better (e.g. both had four spikes), but the spike-times were still off by several milliseconds (spike-time difference 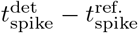 of the last two spikes: 8.2 ms, 3.9 ms). The probabilistic solver revealed this numerical uncertainty (SD of the spike-time *t*_spike_ of the four spikes: 3.7 ms, 5.0 ms, 6.6 ms, 4.1 ms).

Similarly, for the “Inhibition-induced bursting” neuron the solution from the original solver and the deterministic FE solver were qualitatively broadly consistent with the reference solution (Fig. 2C). In all simulations, the neuron responded with spike bursts to a negative stimulus current step. The spike-times and the number of spikes of the original solution (*n*_spikes_ = 11) and the deterministic FE solution (*n*_spikes_ = 14) differed substantially from the reference (*n*_spikes_ = 33) though, with the FE solution having only two bursts instead of three during the simulated period. Here, the probabilistic solver revealed the substantial uncertainty in the spike-times and number of spikes 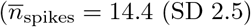, where 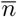 denotes the sample mean), with more than half of the samples having a third burst (Fig. 2C, bottom). All 16 simulated parametrizations are shown in Fig. S1.

To provide an example of a neuronal network, we simulated the STG model for two parametrizations (Figs. 2D and 2E) that only differ in their synaptic conductances (see Section 2.5.3). We computed solutions with a reference solver, a deterministic and a probabilistic EE solver. We focused the analysis on the LP neuron for simplicity. For the first parametrization (Fig. 2D), the LP neuron showed continuous spiking in all simulations. Similar to the HH neuron, we found differences in the exact spike-times and number of spikes between the reference (*n*_spikes_ = 17) and the deterministic EE solution (*n*_spikes_ = 13). The uncertainty was again revealed by the probabilistic solver 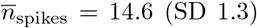; Fig. 2D). The second parametrization resulted in a different spiking behavior of the LP neuron. Here, the neuron started to fire at a high frequency for a prolonged time after approximately two seconds. In the reference solution, the neuron continued to fire. In contrast, in the deterministic solution, the neuron stopped after about another two seconds to then start another burst shortly later. While this also happened in all generated samples from the probabilistic solvers, the sample distribution still indicated a high uncertainty about the duration of the firing periods (Fig. 2C). Simulations of all five synaptic parametrizations from the original paper [25] are shown in Fig. S2.

Finally, we turned to a single STG neuron and stimulated the response to a step stimulus (Fig. 3) based on the original publication [35]. Here, we compared the numerical uncertainty in two different state variables, namely the voltage *v*(*t*) (Fig. 3A) and the intracellular calcium Ca(*t*) (Fig. 3B). We found that the numerical uncertainty differed strongly between these state variables, and was much higher for *v*(*t*) (Fig. 3). While this is expected because of the transient and brief nature of spikes in contrast to the slower changing calcium, it highlights the power of probabilistic ODE solvers, as they can guide the choice of the solver and step-size parameter dependent on the quantity of interest and the desired accuracy without requiring detailed knowledge about the model and its kinetics.

**Figure 3:**
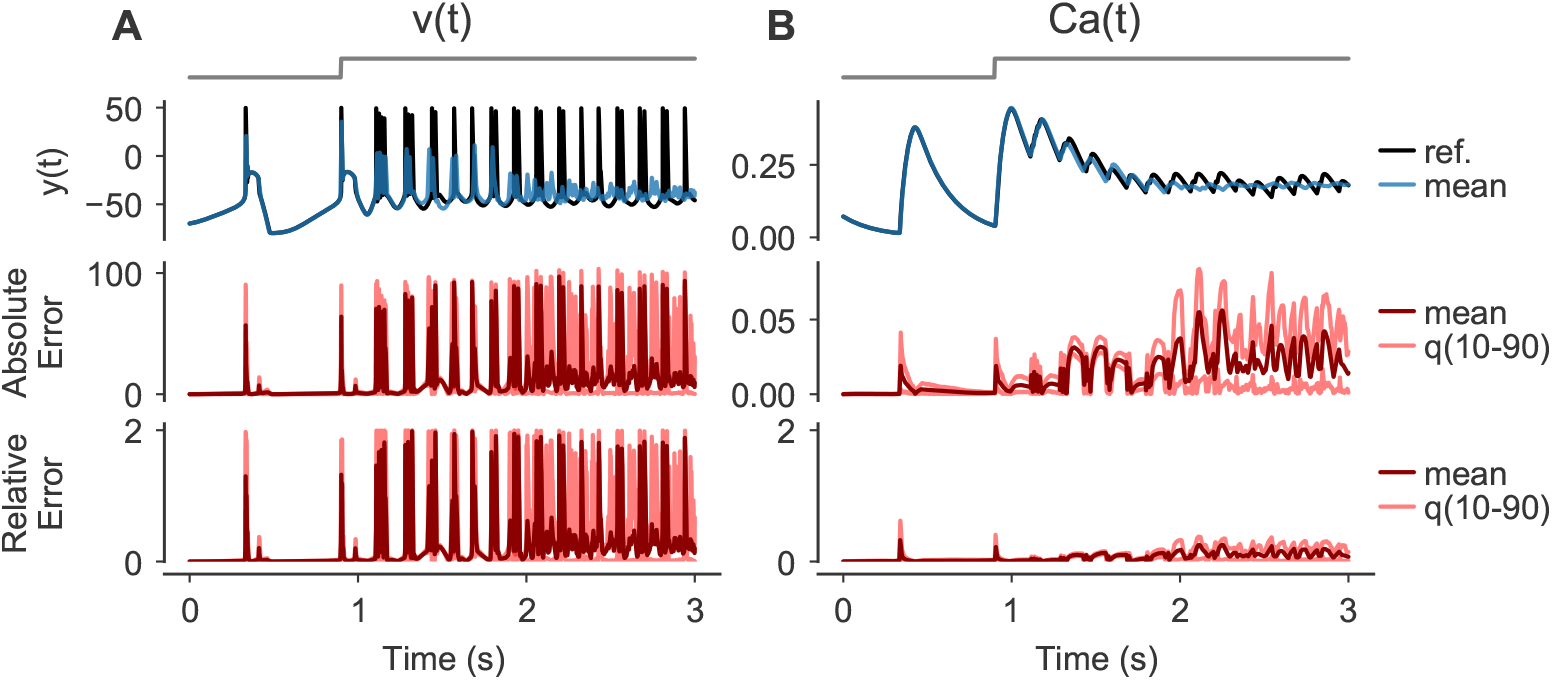
Numerical uncertainty can vary between state variables. **(A, B)** Simulations of a single STG neuron in response to a step-stimulus (gray, top row). Solutions were computed using a reference solver and by drawing 100 samples from a 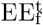 solver with Δ*t*=0.1 ms. The reference solution (black) and the mean over the samples (blue) is shown for two state variables: the membrane potential *v*(*t*) (A, second row) and the intracellular calcium Ca(*t*) (B, second row). For both state variables, the absolute error AE(*t*) = |*y*_sample_(*t*) − *y*_ref._(*t*)| (third row) and the relative error RE(*t*) = |*y*_sample_(*t*) − *y*_ref._(*t*)|*/* max(|*y*_sample_(*t*)|, |*y*_ref._(*t*)|) (bottom row) between sample and reference traces are shown as means and the 10th and 90th percentiles over all samples, respectively.

All the examples in Fig. 2 used first order methods. To also provide an example where higher order solvers with low tolerances yield solutions qualitatively different from the reference solution, we simulated the classical HH neuron’s response for 50 ms to a step stimulus with an amplitude of 0.022 406 mA and *t*_onset_ = 10 ms and *t*_offset_ = 40 ms. This amplitude did not evoke a single spike in the reference solver (Fig. 4A), but was very close to the threshold, i.e. slightly larger amplitudes (e.g. 0.022 410 mA) did produce a spike for the reference solver. When simulating this model with a RKDP_a_ solver, we found that for tolerances of *κ*=1e−3 and *κ*=1e−5 the solutions did contain a spike (Figs. 4A and 4B). Only a tolerance as small as *κ*=1e−7 yielded a solution with no spike for this solver (Fig. 4C). Simulating the model with probabilistic solvers revealed this numerical uncertainty for both *κ*=1e−3 and *κ*=1e−5, with a fraction of samples containing one and a fraction containing zero spikes in both cases (Fig. 4D).

**Figure 4:**
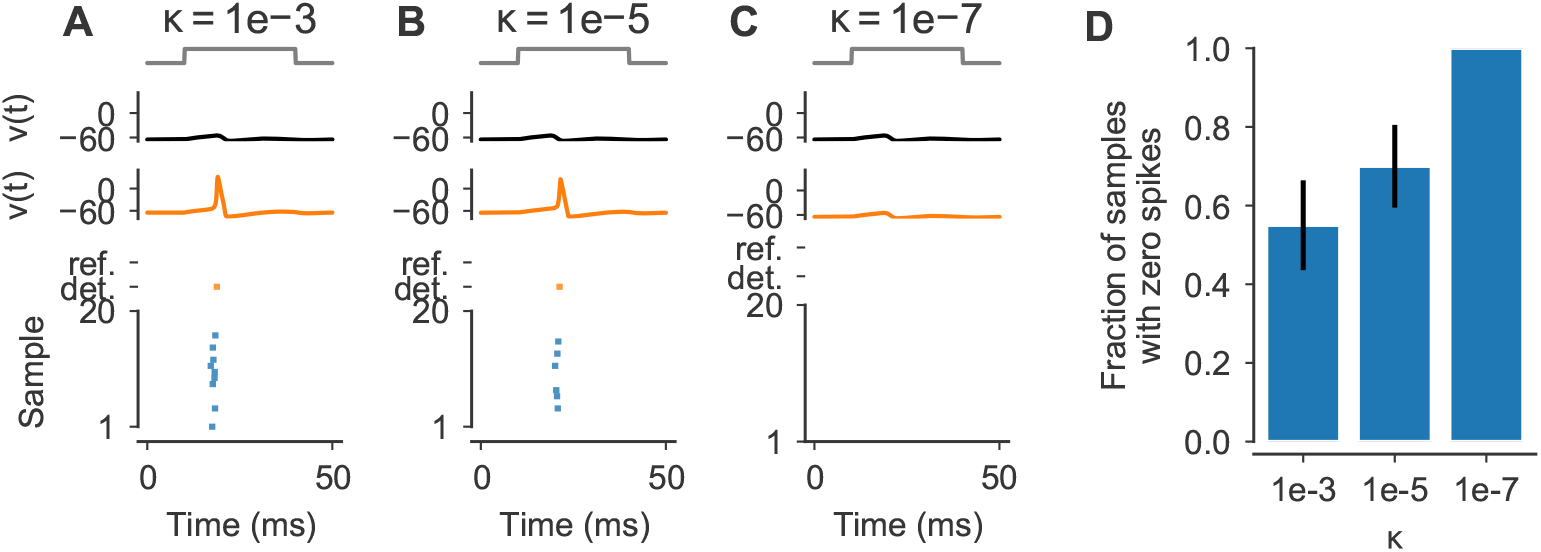
Numerical uncertainty affects also higher order methods. **(A-C)** Simulations of the classical HH model for the step stimulus *I*_Stim_ with an amplitude of 0.022 523 5 mA (normalized stimulus in gray). Solutions for *v*(*t*) are shown for a reference solver (black) and a deterministic RKDP_a_ solver with *κ*=1e−3, *κ*=1e−5 and *κ*=1e−7, respectively (orange). *Bottom panels*: Spike-times of the reference (black), the deterministic solutions (orange) and for 20 samples from probabilistic 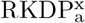 solvers with *κ*=1e−3, *κ*=1e−5 and *κ*=1e−7, respectively (blue). **(D)** Fraction of samples (*n* = 20) from the probabilistic solvers in (A-C) that had no spike, shown as mean and standard error. All other samples had exactly one spike.

### 3.2 Probabilistic solvers can guide solver selection

To demonstrate how probabilistic ODE solvers can be used to compare the accuracy vs. run time tradeoff between different solver schemes, we simulated the HH neuron’s response to the noisy step stimulus (Fig. 5A) using the following probabilistic solvers: 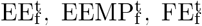 and 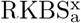. To this end, we computed the Mean Absolute Error between voltage trace samples *v*(*t*) and the reference (MAE_SR_) for each solver as an estimate of the numerical error induced. We compared these errors to the number of ODE evaluations a corresponding deterministic solver would need. We found that the exponential integrators EE and EEMP allowed computing solutions with the fewest ODE evaluations, as they terminated successfully even for the relatively large step-size Δ*t*=0.5 ms (Fig. 5B). In contrast, when using the FE solver, all step-sizes Δ*t* ≫ 0.05 ms resulted in floating-point overflow errors and therefore in both useless and incomplete solutions. However, when choosing a sufficiently small step-size of Δ*t* ≤ 0.05 ms the samples obtained with the FE method had on average a smaller error compared to the EE method (Fig. 5B). From the methods tested, the adaptive RKBS method was the most efficient one, i.e. it produced the most accurate solutions for the fewest number of ODE solutions, but it also required a substantially higher number of minimum ODE evaluations to successfully terminate compared to the exponential integrators (Fig. 5B).

**Figure 5:**
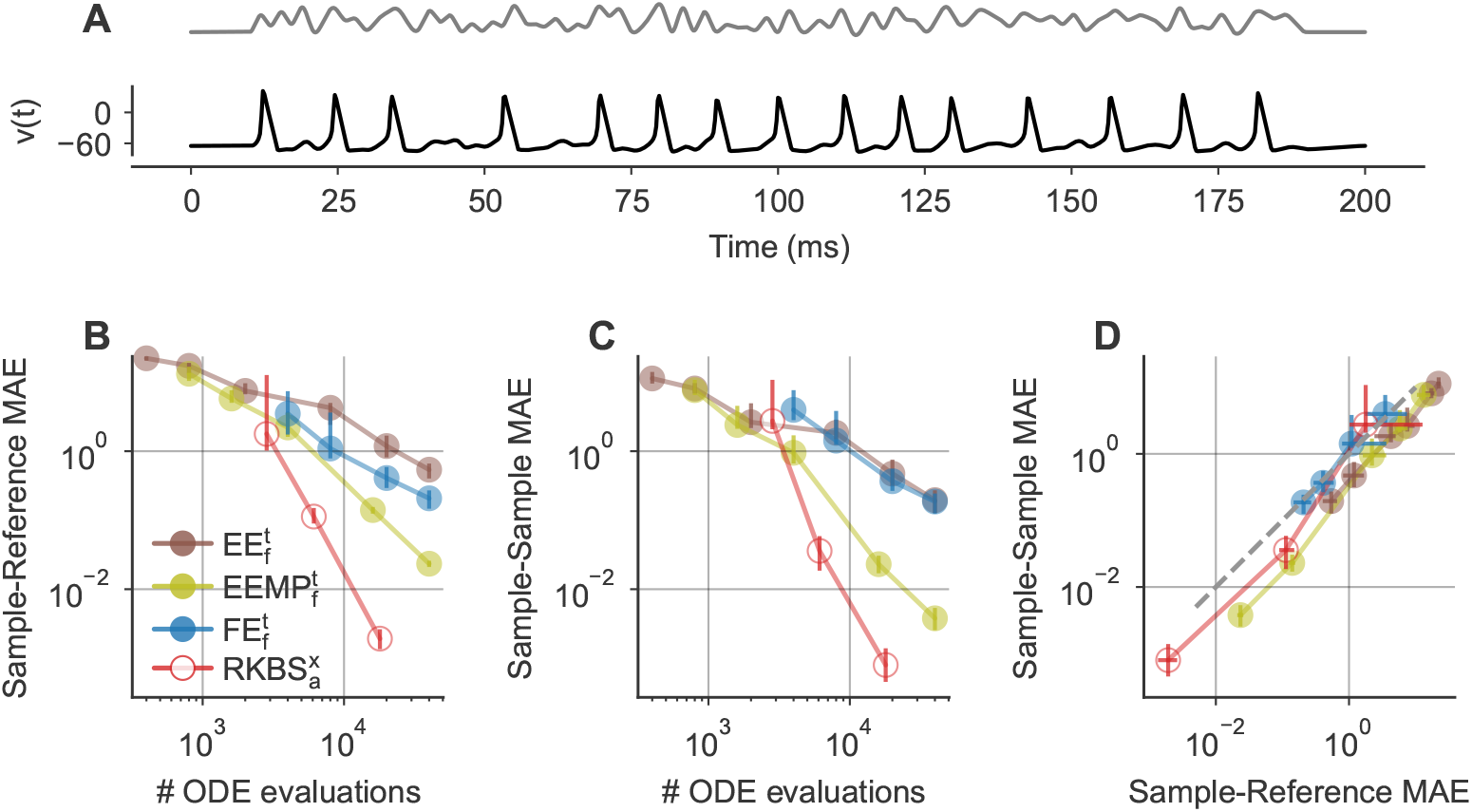
Probabilistic solvers can be used to compare different solver schemes. **(A)** Reference solutions of *v*(*t*) (black) for the Hodgkin-Huxley model stimulated with the noisy step stimulus (normalized stimulus in gray). **(B)** Mean Absolute Errors MAE_SR_ between sample traces of *v*(*t*) and the respective reference solutions for different solver schemes (legend) and step-sizes / tolerances. Mean Absolute Errors from 100 samples are shown as medians (dots) and 10th to 90th percentiles (vertical lines) as a function of the number of ODE evaluations of a corresponding deterministic solver (x-axis). **(C)** As in (B), but for sample-sample Mean Absolute Errors MAE_SM_. **(D)** As in (B), but for sample-sample Mean Absolute Errors MAE_SM_ vs. sample-reference Mean Absolute Errors MAE_SR_.

In principle, a very similar analysis could also have been done with deterministic solvers. However, probabilistic solvers have two advantages. First, they yield sample distributions instead of single solutions which make it possible to compute confidence intervals etc. when comparing different solver outputs. Second, and more crucially, probabilistic solvers do not require a reference solution to estimate the numerical error in a solution. For a sufficiently calibrated probabilistic solver, the sample distribution, i.e. the solver’s output, can be used to estimate the numerical error of the solver. In Fig. 5C we computed the sample-sample distances MAE_SM_, which are independent of the reference, for the same samples used in Fig. 5B. We found that the mean sample-reference distances 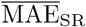 were highly similar to the respective mean sample-sample distances 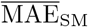 for all solvers (Figs. 5B–5D). Therefore, the solver comparison described above could have also been based on MAE_SM_ instead of MAE_SR_, and therefore would not have required a reference solution.

### 3.3 Calibration of probabilistic solvers

The mean sample-sample distance 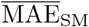 is only then a good approximation to the mean sample-reference distance 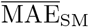 (as for example in Fig. 5) when the probabilistic solver is well calibrated. Ideally, the magnitude of the perturbation is large enough to capture the numerical uncertainty of the underlying numerical integration, but it is not too large to severely reduce the accuracy of the integration scheme. To quantify the calibration of different solvers, we therefore defined two metrics, the ratio 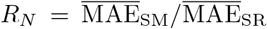 and the ratio 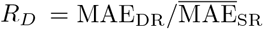, where MAE_DR_ is the distance between a corresponding deterministic solution and the reference. *R*_*N*_ is close to zero if the perturbation is too small (i.e. the sample-sample distance is much smaller than the sample-reference distance) and close to one if the perturbation is sufficiently large to not underestimate the numerical uncertainty (i.e. the sample-sample distance can be used as an approximate measure of the sample-reference distance). *R*_*D*_ is close to one if the perturbation is either too small to affect the model output (i.e. all samples are approximately equal to the deterministic solution) or if samples are on average approximately equally close to the reference than the deterministic solution. *R*_*D*_ converges to zero, if the perturbation is too large and the perturbation severely reduces the solver accuracy. Note that *R*_*D*_ can also take values larger than one, which happens when the perturbation increases the solver accuracy on average (e.g. see Fig. 2C). This happens for example when the deterministic solution is missing a spike but almost reaches the model’s spike threshold, and the perturbation is strong enough to generate the missing spike in some samples.

For a well calibrated solver, *R*_*N*_ is close to one, such that the sample-reference distance 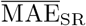 can be estimated from sample-sample distance 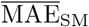 while *R*_*D*_ is close to or larger than one, such that the perturbation does not decrease the solver accuracy.

The magnitude of the perturbation can be adjusted with the perturbation parameter *σ* that we defined for both the state and step-size perturbation (see Section 2.1). To analyze how the parameter *σ* affects the calibration of the perturbation and to test for which *σ* the solvers are well calibrated, we simulated the classical HH neuron in response to the noisy step stimulus with probabilistic solvers for a range of perturbation parameters (Fig. 6). First, we used a probabilistic 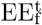 solver and computed MAE_SM_, MAE_SR_ and MAE_DR_ and the ratios of the distribution means *R*_*N*_ and *R*_*D*_ for five different *σ* ranging from 0.0625 to 16 (Figs. 6A–6C). We found that with increasing *σ*, the mean sample-sample distance 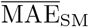 converged to the mean sample-reference distance 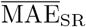, and for sufficiently large *σ* the mean sample-sample distance 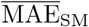 could therefore be used as an approximate measure of mean sample-reference distance 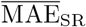 (Figs. 6A and 6B). For example, for *σ*=0.25, the perturbation magnitude was too small and the solver was underestimating the numerical uncertainty: Here, the mean sample-sample distances was much smaller 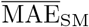 (0.33) than mean the sample-reference distance 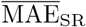 (4.35) (Fig. 6A) with all sample-reference distances distributed narrowly (MAE_SR_ 10th to 90th percentiles: [4.14, 4.56]) around the sample-deterministic distance MAE_DR_ (4.36), indicating that all samples were very close to the deterministic solution, despite the numerical error. When using *σ* ≥ 4, the mean sample-reference distance was higher than the deterministic-reference distance (Figs. 6A and 6C), indicating a loss of solver accuracy caused by the perturbation (e.g. *R*_*D*_ = 0.51 for *σ*=8). The best calibration was achieved with 1 ≤ *σ* ≤ 4, with distributions of MAE_SM_ close to MAE_SR_ (*R*_*N*_ : 0.51, 0.83 and 0.97 for *σ*=1, *σ*=2 and *σ*=4, respectively; Figs. 6A and 6B) and with the mean sample accuracy close to the accuracy of the deterministic solution (*R*_*D*_: 1.11, 1.08 and 0.74 for *σ*=1, *σ*=2 and *σ*=4, respectively; Figs. 6A and 6C).

**Figure 6:**
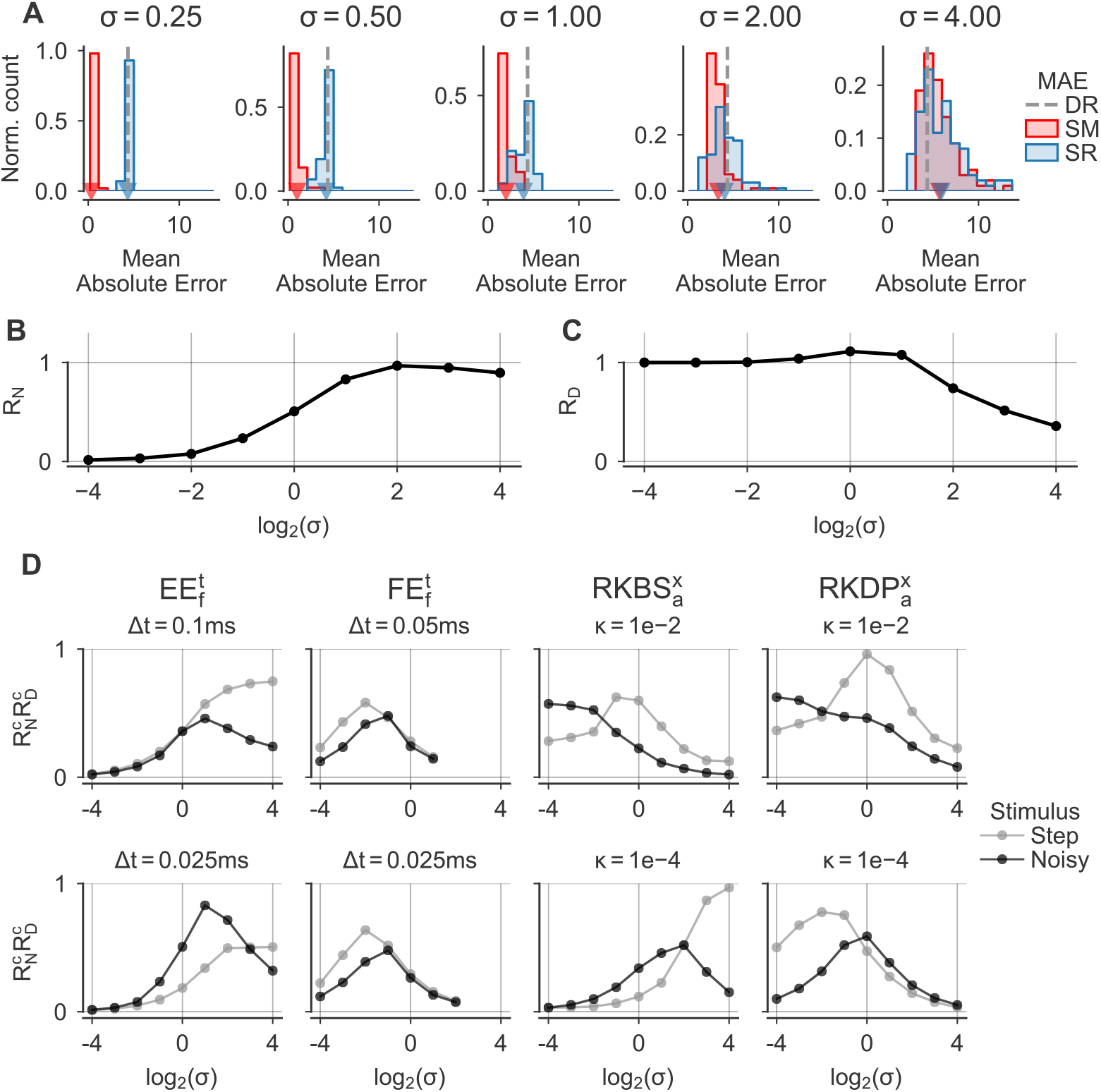
Default calibration of probabilistic solvers is good but not optimal. Simulations of the Hodgkin-Huxley model with different probabilistic solvers and perturbation parameters *σ* ranging from 0.0625 to 16. **(A)** Distributions of the Mean Absolute Error between voltage traces *v*(*t*) computed with a probabilistic 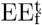 solver with Δ*t*=0.025 ms, a deterministic version of this solver, and a reference solver for five perturbation parameters (titles). Solutions were computed for the noisy step stimulus. In each panel: Mean Absolute Error distributions were computed between 100 samples and the reference as MAE_SR_ (blue histograms), between the samples and the sample mean MAE_SM_ (red histograms) and between the deterministic and reference solution as MAE_DR_ (dashed grey line). Means of the distributions are highlighted (triangles). **(B)** Ratios of the Mean Absolute Error distribution means 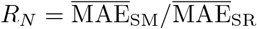 as a function of the perturbation parameter. **(C)** As in (B), but for ratio 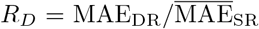. **(D)** Clipped ratio products 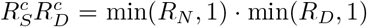 for different solvers (column titles) and step-sizes Δ*t* / tolerances *κ* (panel titles) for the step (grey) and the noisy step (black) stimulus as a function of the perturbation parameter *σ* (x-axis).

To provide an overview of the calibration for different solvers settings, we defined the clipped ratio product 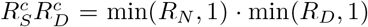, which is ideally close to one. We used clipped values, because in some cases *R*_*S*_ and *R*_*D*_ took values larger than one which makes their product more difficult to interpret (Fig. S3). 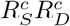 is close to zero for either an underestimation of the numerical uncertainty 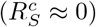 or for a too strong perturbation that renders the solver output useless 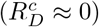.

We computed 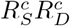 for different probabilistic solvers and step-sizes—including the 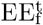 solver with Δ*t*=0.025 ms used above—for the HH neuron stimulated with the step and noisy step stimulus (Fig. 6D). The respective values for *R*_*N*_ and *R*_*D*_ are shown in Fig. S3. We found that the default perturbation parameter *σ* produced reasonably calibrated solutions in all cases. However, in most cases *σ*=1 was also not ideal. For 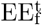, larger values (e.g. *σ*=2 or *σ*=4) resulted in better calibration, whereas for 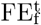 the calibration was improved using smaller values (e.g. *σ*=0.5 or *σ*=0.25). The adaptive Runge-Kutta methods RKBS and RKDP were well calibrated for a wide range of perturbation parameters *σ*, including very small ones (e.g. *σ*=0.0625), especially in the high tolerance case (*κ*=1*e*−2). This is likely because even small perturbations cause the solvers to take different step-sizes and therefore to evaluate the ODE at different time points.

### 3.4 Computational overhead

Probabilistic solvers based on state or step-size perturbation increase the computational costs for two reasons. First, they are sampling based and require computing multiple solutions for a single IVP. While this process can be parallelized, it may nevertheless come with a computational overhead, especially if it conflicts with other computations using parallelized model evaluation, e.g. in simulation based inference where the same model is evaluated for different model parameters [37, 38]. Second, probabilistic solvers induce a computational overhead per solution computed relative to their deterministic counterparts. We analyzed both aspects in the following.

#### 3.4.1 Required number of samples

To empirically determine the number of samples necessary to obtain a reliable measure of numerical uncertainty, we simulated the classical HH neuron with probabilistic solvers for the step and noisy step stimulus. To this end, we computed mean sample-sample distances 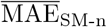 for small numbers of samples *n*, and divided them by the mean sample-sample distances for a much larger number of samples (300) to obtain normalized sample-sample distances 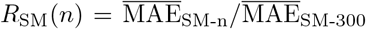 (Figs. 7A and 7B). We found that often already two samples were sufficient to get a good estimate of the sample-sample distance. e.g. for the step stimulus and *n* = 2, more than half of the *R*_SM_ were in [0.67, 1.5] with little difference between the solvers 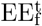 (Fig. 7A) and 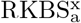(Fig. 7B).

**Figure 7:**
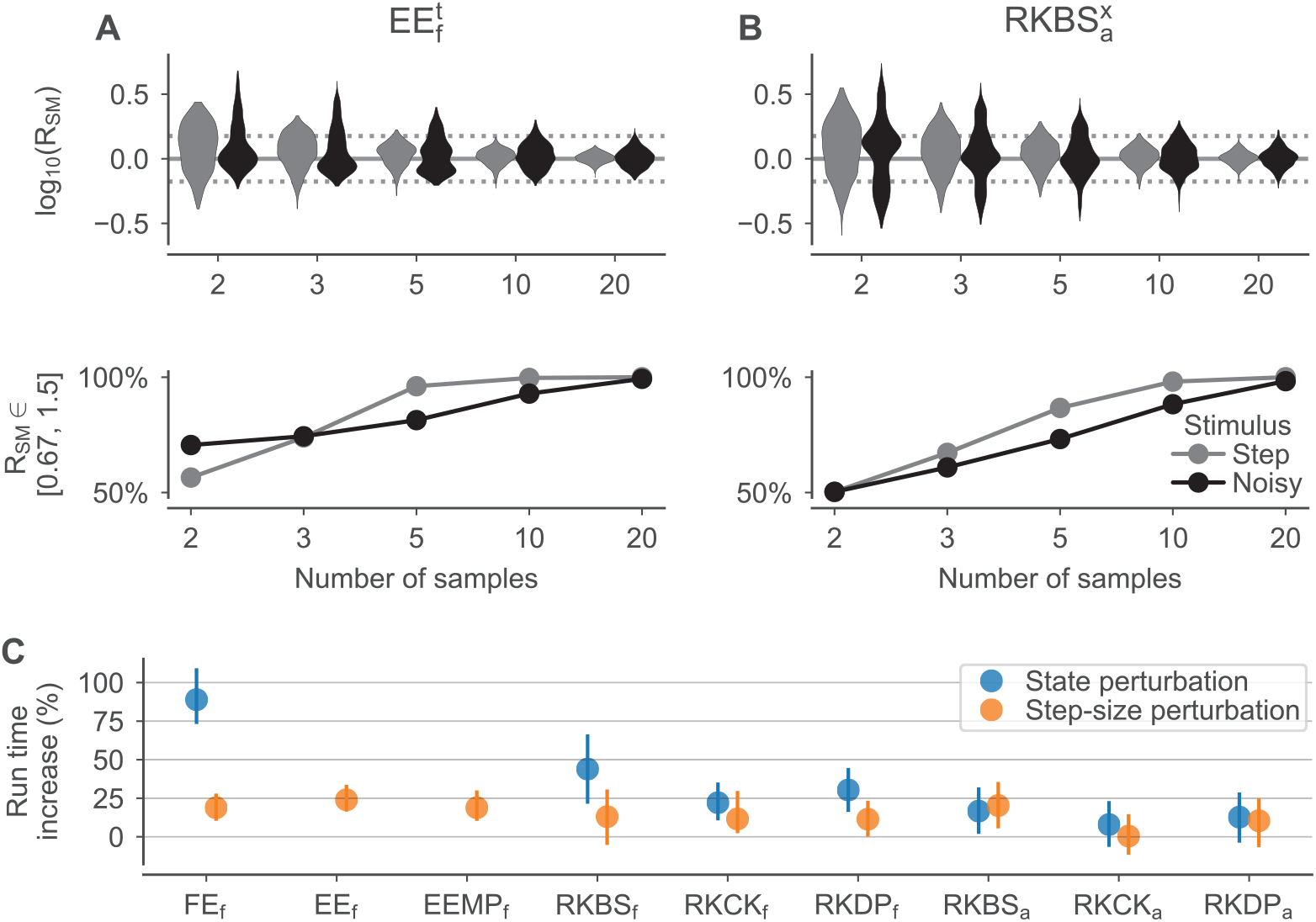
The computational overhead of probabilistic solver is moderate. **(A**,**B)** Bootstrapped distributions of normalized sample-sample distances 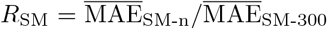 as a function of the numbers of samples *n. R*_SM-n_ were computed for the classical HH neuron simulated with a 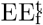 with Δ*t*=0.1 ms (A) and a 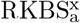 solver with *κ*=1e−2 (B) for the step (grey) and noisy step (black) stimulus, respectively. *Top:* Distributions over log_10_(*R*_SM_) computed by 1000 times repeated random sub-sampling of 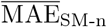 from all 300 generated samples. Vertical lines highlight *R*_SM_ values of 0.67, 1 and 1.5. *Bottom:* Percentages of the bootstrapped *R*_SM_ distributions in the interval [0.67, 1.5]. **(C)** Relative run times for different solver schemes measured for the classical HH neuron with the noisy step stimulus. For every solver, 100 samples were simulated for both a probabilistic and a corresponding deterministic solver. Relative run times were computed by dividing the run times of the probabilistic samples by the run times of the respective deterministic samples. Distributions were computed by bootstrapping 10000 ratios and are shown as medians and the 10th to 90th percentiles. The step-size of fixed step-size methods was Δ*t* = 0.05 ms and the tolerance of adaptive methods was *κ*=1e−4. For EE and EEMP, only the step-size perturbation was used.

#### 3.4.2 Overhead per sample

In addition to in the computational overhead caused by the computation of multiple samples, probabilistic methods also come with a computational overhead per solution. For the state perturbation [18] this overhead has three components. First, one needs to compute the local error estimator, which only causes overhead for fixed step-sizes since for adaptive methods the local error estimator needs to be computed anyway. The second potential source of overhead is that the “First Same As Last” property—i.e. that the last stage in one step can be used as the first stage of the next step, which is used in RKBS and RKDP—is not applicable. This is because the last stage is computed before the perturbation, and after the perturbation the evaluation of the ODE is not valid anymore. Lastly, the perturbation itself, which includes sampling form a Gaussian, needs to be computed.

In total, this overhead is relatively small for higher order methods optimized for step-size adaptation like RKBS, RKCK and RKDP. For example, the state perturbation for RKDP_a_ increases the number of ODE evaluations per step from six to seven (+16%) due to the loss of the First Same As Last property, and for RKCK_a_—which does not make use of this property—no additional ODE evaluation is required. However, for first order methods like FE this overhead severely reduces the computational efficiency because instead of a single ODE evaluation per step, a state perturbed version needs two (+100%). Additionally, lower order methods typically require more steps in total compared to higher order methods, because they are typically used in combination with smaller step-sizes. This increases the total computational costs of the perturbation itself, which is done once per step. For the step-size perturbation, the overhead is reduced to the perturbation and, for adaptive step-size methods, the loss of the First Same As Last property.

To quantify this overhead empirically, we simulated the HH neuron with different probabilistic solvers and their deterministic counterparts and compared the run times relative to each other. As expected, for the state perturbation, the computational overhead was larger for the lower order methods (Fig. 7C; on average for 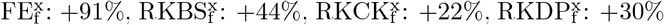 The adaptive methods—where the local error estimates were computed not only for the probabilistic, but also for the deterministic methods—showed the smallest increase in run times (+12% on average across all adaptive methods), with 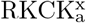, not using the First Same As Last property, having the least overhead (+8%). For the step-size perturbation, the increase in run times was on average smaller (+15% on average across all methods) and without large differences between the solver schemes and the usage of adaptive or fixed step-sizes.

### 3.5 Limitations

Finally, we turned back to the “DAP” IN model, to illustrate limitations of the approach. For this neuron model we had found a large difference in the number of spikes for the fixed step-size methods, like the original solver, compared to the reference solution (Fig. 8A). While the reference solution had eight spikes during the simulated period, the original solution had only one and a deterministic FE solver had two. While the probabilistic solver 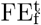 arguably indicated some numerical uncertainty 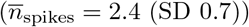, the number of spikes was still much lower compared to the reference. To better understand the source of this numerical uncertainty, we simulated the “DAP” neuron model with different probabilistic solvers, 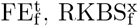 and 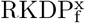.

**Figure 8:**
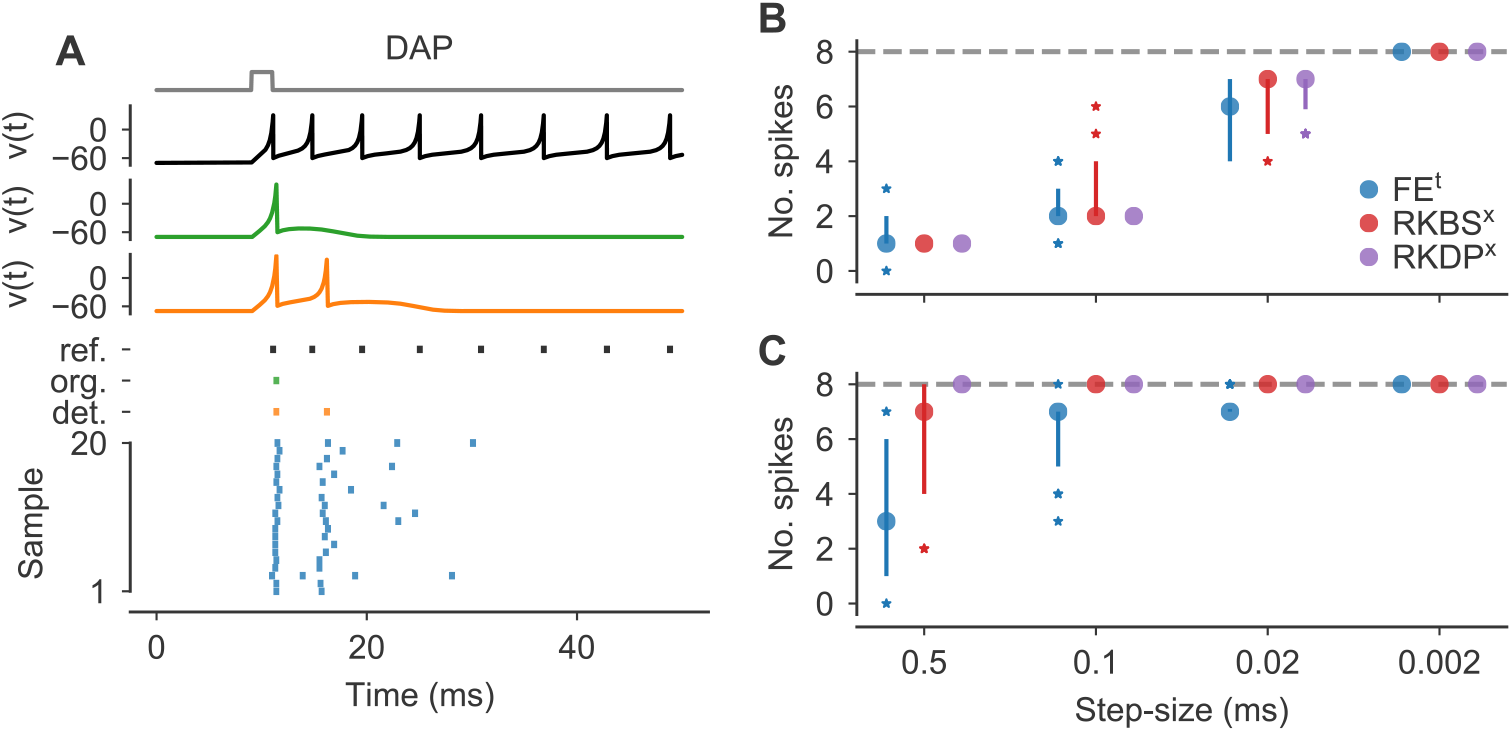
Probabilistic solver only account for errors in numerical integration. **(A)** Simulations of the “DAP” IN model with a pulse stimulus *I*_Stim_ (normalized stimuli in gray). Solutions for *v*(*t*) are shown for a reference solver (black), the original solver scheme (green) and a deterministic FE solver (orange). Based on the original publication, the step-size Δ*t* was set to 0.1 ms. For plotting, *v*(*t*) were clipped at 30. *Bottom panel* : Spike-times are shown for the reference (black), the original solver solution (green), the deterministic FE solution (orange) and for 20 samples from a probabilistic 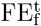 solver (blue). Samples were sorted by the number of spikes. **(B**,**C)** Number of spikes for the “DAP” IN model (see Fig. 2C) dependent on the step-size (x-axis) for different solver methods (legend) for fixed (**A**) and pseudo-fixed (**B**) step-sizes, respectively. Number of spikes are shown for 40 samples as medians, 10th to 90th percentiles (vertical lines) and outliers (stars). The dashed horizontal line refers to the reference solution (see Fig. 2C).

First, we simulated the neuron for different fixed step-sizes. We found that all probabilistic solvers underestimated the true number of spikes when using relatively large fixed step-sizes (Fig. 8B). For the largest step-size tested, Δ*t*=0.5 ms, only the 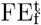 solver indicated uncertainty in the number of spikes, whereas for 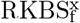 and 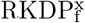 all samples had only a single spike. When using smaller step-sizes, the probabilistic solvers’ outputs were more indicative of the numerical uncertainty. For Δ*t*=0.02 ms, all solvers produced outputs that were closer (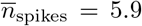 for 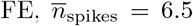 for RKBS and 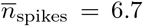 for RKDP) to the reference solution and all methods indicated uncertainty in the number of spikes. With the very small step-size Δ*t*=0.002 ms, all samples from all solvers showed the same number of spikes as the reference and the probabilistic solvers indicated no remaining uncertainty about the number of spikes here.

While these results may be unsatisfactory at first glance, they are not necessarily unexpected. The probabilistic solvers used here can only capture the uncertainty arising through the numerical integration; they can not capture the error that is introduced by restricting spikes to occur only on a fixed time grid, which is the case for the fixed step-size solvers. We therefore simulated the neuron for the same solvers and step-sizes again, but allowed the solver to take intermediate steps (see Eq. (10)) every time a reset occurred. When using these “pseudo-fixed” step-sizes, we found that 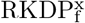 still did not indicate uncertainty in the number of spikes for any step-size tested, but now all samples had the same number of spikes as the reference (Fig. 8C). And while 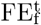 and 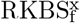 still underestimated the number of spikes for larger step-sizes on average (e.g. for Δ*t*=0.5 ms: 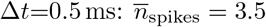 for FE and 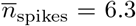 for RKBS), both indicated high numerical uncertainty (e.g. for Δ*t*=0.5 ms: *q*_90_(*n*_spikes_) = 6 for FE and *q*_90_(*n*_spikes_) = 8 for RKBS, where *q*_90_ is the 90th percentile).

## 4 Discussion

The outcome of neuron simulations is affected by numerical uncertainty arising from the inevitably finite step-sizes used in numerical ODE integration. With standard solvers there is no straightforward way to quantify how this uncertainty affects quantities of interest such as spike-times and the number of spikes.

In this study, we demonstrated how probabilistic solvers can be used to quantify and reveal numerical uncertainty in commonly used neuron models. Crucially, these solvers can be easily implemented and do not require a detailed understanding of the underlying kinetics of the neuron model of interest.

Further, we showed that numerical uncertainty can affect the precise timing and sometimes even the number of the spikes in simulations of neuron models commonly used in neuroscience. We also found that some models and parametrizations are more susceptible to numerical uncertainty than others, and that some solvers employed in the neuroscience literature yield rather large uncertainties. These findings highlight the need for a thorough quantification of numerical uncertainty in neuroscience simulations to strike an informed balance between simulation time and tolerated uncertainty.

The idea to quantify the accuracy or numerical errors of different solvers for mechanistic models in neuroscience is not new. For example, Butera and McCarthy [39] showed that for small step-sizes, the forward Euler method produces more accurate solutions than the exponential Euler method, which is in agreement with our findings. Börgers and Nectow [11] on the other hand argued that for Hodgkin-Huxley-like systems exponential integrators— such as exponential Euler and the exponential midpoint Euler—are often the best choice, as they allow for much larger step-sizes especially when high accuracy is not necessary, which is again what we observed. Stewart and Bair [10] argued in favor of the Parker-Sochacki integration method and showed that it can be used to generate highly accurate solutions for both the Izhikevich and Hodgkin-Huxley model. However, this method has the disadvantage that the ODE system at hand has to be put into the proper form and therefore requires specific knowledge about the model and solver. In a more recent study, Chen et al. [12] recommended to use splitting methods, such as second-order Strang splitting, instead of exponential integrators.

In contrast to these studies, probabilistic solvers offer a more general approach to tackle the problem of numerical uncertainty. Instead of finding the “best” solver for a specific problem, they produce an easy-to-interpret uncertainty measure that can be analyzed without specific knowledge about the solver or solved neuron model. This allows to easily assess if a solver is sufficiently accurate for a given research question. It can therefore facilitate both the choice of the solver and choice of solver settings such as the step-size.

In this study, we used two simple probabilistic solvers that build on deterministic solver and stochastically perturb the numerical integration. For both, the state [18] and the step-size perturbation [22] method, it is crucial that the perturbation is of the right order to neither underestimate the numerical uncertainty nor to reduce the solver accuracy unnecessarily. To be able to adjust the perturbation we introduced a perturbation parameter for both the state and step-size perturbation. We found that using the default value for this parameter yielded good calibration in most cases, but slight adjustments often improved the calibration further.

The state and step-size perturbation are conceptually quite similar, but the two methods have some clear advantages and disadvantages with respect to each other. The step-size perturbation requires a local error estimator to be calibrated. This is disadvantageous because it requires a method for local error estimation which can introduce a relatively large computational overhead per solution for lower order methods like FE. The step-size perturbation may therefore a better choice for lower order methods. However, for higher order methods like RKDP this difference vanishes and both approaches require an equally small computational overhead per solution. Another advantage of the step-size perturbation is that it preserves desirable properties of the underlying solver schemes [22]. For example, when Hodgkin-Huxley-like models are solved with exponential integrators like EE or EEMP, the state variables of the activation and inactivation can not leave their domain [0, 1] by design of the solvers, a property preserved by the step-size but not the state perturbation. A downside of the step-size perturbation is that the calibration can be slightly more challenging because the perturbation is influenced by linear scaling of the simulated time, which happens for example if the time unit of the model is changed.

Beyond the two perturbation methods used and discussed in this study, there are probabilistic ODE solvers constructed using techniques from (nonlinear) Gaussian filtering and smoothing [15, 16, 40]. These methods have the advantage that instead of repeatedly integrating the initial value problem, they only require a single forward integration and return local uncertainty estimates that are proportional to the local truncation error. The disadvantage of Gaussian ODE filters and smoothers is that the uncertainty estimates are Gaussian. This restriction can be lifted by replacing Gaussian filters and smoothers with particle filters and smoothers [16]. In particular for large neural network simulations, such efficient methods will be key in quantifying uncertainty.

To further extent the applicability of probabilistic solvers in neuroscience, it will be crucial to also develop and test implicit probabilistic solvers for neuron models. For example, the ODEs of multi-compartment neuron models are typically stiff which makes implicit solvers the better choice for such models [41]. A priori, it is often not easy to judge whether a ODE system is stiff or not. A noteworthy attempt to tackle this problem is the algorithm by Blundell et al. [42] that automatically determines whether an implicit or an explicit solver should be used.

## 5 Acknowledgements

This research was funded by the Deutsche Forschungsgemeinschaft through a Heisenberg Professorship (BE5601/8-1, PB), the Excellence Cluster 2064 “Machine Learning — New Perspectives for Science” (ref number 390727645, PB and PH), ADIMEM (01IS18052C and 01IS18052B to PB and PH) and the Tübingen AI Center (FKZ: 01IS18039A, PB and PH). NK and PH gratefully acknowledge financial support by the German Federal Ministry of Education and Research (BMBF) through Project ADIMEM (FKZ 01IS18052B), as well as by the European Research Council through ERC StG Action 757275 / PANAMA, and funds from the Ministry of Science, Research and Arts of the State of Baden-Württemberg. The authors thank the International Max Planck Research School for Intelligent Systems (IMPRS-IS) for supporting Nicholas Krämer.

## Appendix

### A Local error estimation and step-size adaptation

To compute the local error estimator ***ε***_*t*_ for a single integration step, step solutions were computed with two different numerical methods that were run in parallel to provide two solutions **x**_*a*_(*t* + Δ*t*) and **x**_*b*_(*t* + Δ*t*) for every step, given *t*, Δ*t* and **x**(*t*) (e.g. see Dormand and Prince [28]). In every step, the local error estimator was computed as:

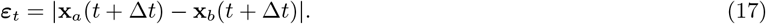

For adaptive step-size methods, the error estimator 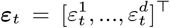 was used to compute an error norm ||**e**|| on **e** = [*e*_1_, …, *e*_*d*_]^*T*^, where *d* was the dimension of the state vector **x**(*t*) = [*x*_1_(*t*), …, *x*_*d*_(*t*)]^*T*^. For every state variable *x*_*i*_, *e*_*i*_ was computed as:

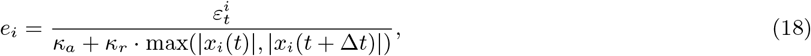

with *κ*_*a*_ and *κ*_*r*_ being the absolute and relative tolerance. For simplicity, we used *κ*_*a*_ = *κ*_*r*_ in all simulations and therefore refer to these parameters as the *tolerance κ*.

||**e**|| was computed as the root-mean-square of **e**, i.e.:

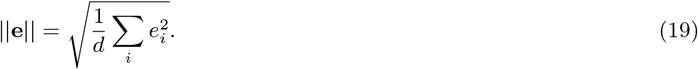

If ||**e**|| *<* 1, the step was accepted, and rejected otherwise. In both cases, the step-size was adapted and the next step-size Δ*t*_next_ was computed as:

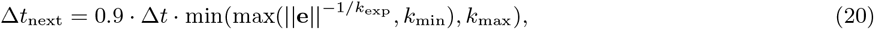

where *k*_min_ and *k*_max_ are the minimum and maximum allowed change factors, that we set to typical values of 0.1 and 5 respectively [9]. *k*_exp_ was 2 for FE, 3 for RKBS, 4 for RKCK, and 5 for RKDP, corresponding to the order of the error estimator. Furthermore, we limited the step-sizes to be always smaller or equal to a maximum step-size Δ*t*_max_, which we set to Δ*t*_max_ = 1 ms for all simulations.

### B Solver details

Runge-Kutta steps were implemented based on the scipy implementation [30]. The Butcher tableau for the RKCK method was taken from [27]. Heun’s method was used as an error estimator for the FE method and implemented as follows. Given *t*, Δ*t*, **x**(*t*), *f* (*t*, **x**(*t*)) and the deterministic FE solution 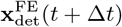 the solution for Heun’s method was computed as:

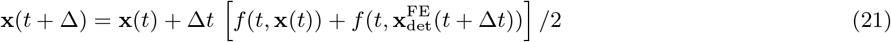

To use the exponential integrators EE and EEMP, the ODEs were cast into the following form:

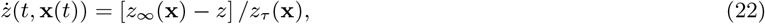

where *z* is a state variable of **x** (e.g. the membrane potential) and *z*_∞_(**x**) and *z*_*τ*_ (**x**) are functions depending on **x** but not explicitly on *t*. For a derivation of these functions for the HH model see for example [43]. The EE step was implemented as:

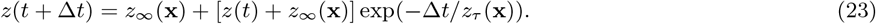

The second order exponential integrator EEMP proposed by Börgers and Nectow [11] builds on the EE method. Given *t*, Δ*t*, **x**(*t*), the half-step EE solution 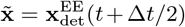 and the evaluations of 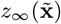 and 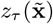 at the half-step, the solution for *z* using the EEMP method was computed as:

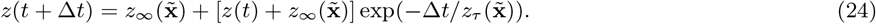

We also implemented the solver used in the original implementation of the IN neurons, where the IVP was solved with a method similar to FE of fixed step-size Δ*t* [23]. The implementation differs from a standard FE scheme in so far, as *v* and *u* are updated subsequently:

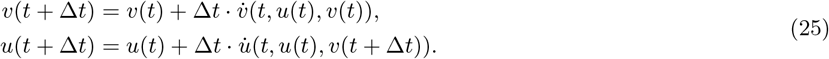

### C Neuron model parameters

The neuron models simulated in this study were parametrized as follows. The parameters ***θ*** = [*a, b, c, d*] and *I*_Stim_ of the IN model and the respective original step-sizes Δ*t* were taken from https://www.izhikevich.org/publications/figure1.m [23].

The three maximum conductances for the classical HH neuron (see Eq. (12)) were set to 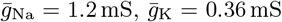 and 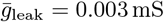. The membrane capacitance was set to *C* = 0.01 μF (see Eq. (11)). For all STG neurons, we set the membrane area to *A* = 0.628 *×* 10^−3^ cm^2^ and the membrane capacitance to *C* = *A* · 1 μF*/*cm^2^ (see Eq. (11)). An STG neuron has eight maximum channel conductances (see Eq. (12)):

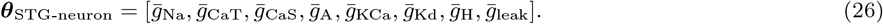

For the single STG neuron, we set ***θ***_STG-neuron_ = *A* · [400, 2.5, 10, 50, 20, 0, 0.04, 0] mS*/*cm^2^, taken from an example in [35]. The STG neuronal network consists of three neuron models ABPD, LP and PY. The network is parametrized by the three neurons’ conductances:

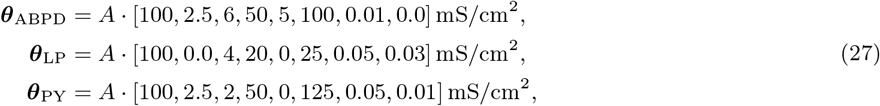

where *A* = 0.628 *×* 10^−3^ cm^2^ and the synaptic conductances ***θ***_syn_:

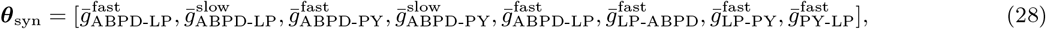

where for example 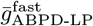 is the maximum conductance of the fast synapse connecting neuron ABPD (presynaptic) to neuron LP (postsynaptic). We simulated the network for five different synaptic parametrizations taken from the original publication [25]:

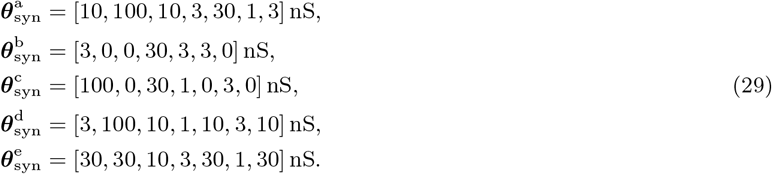

The frequencies *f*_*s*_ (Eq. (15)) for the fast and slow synapses were 25 Hz and 10 Hz, and the reversal potentials *E*_*i*_ (Eq. (14)) were −70 mV and −80 mV, respectively [25].

**Figure S1:**
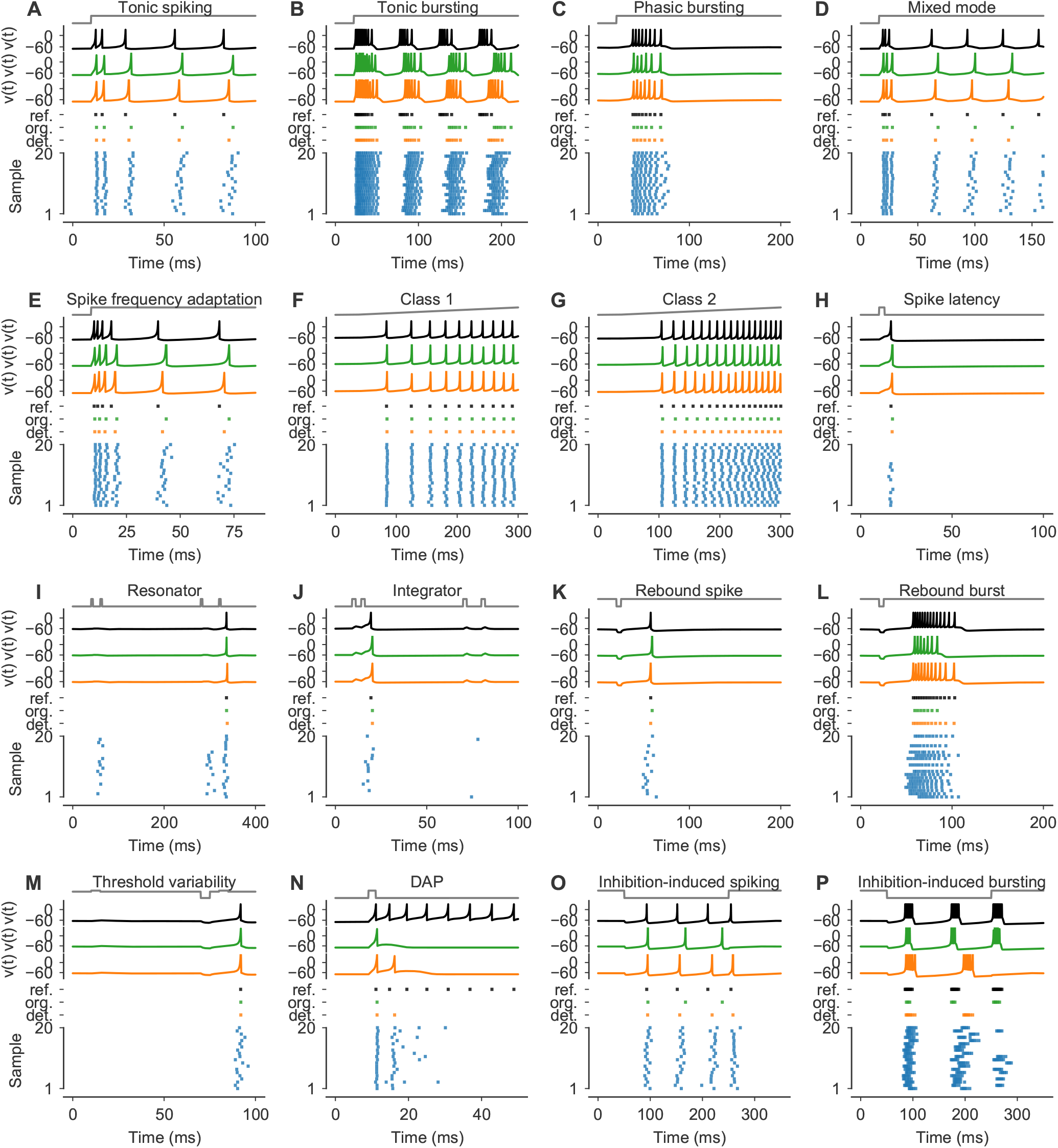
**(A-P)** As in Figs. 2B and 2C. Simulations of the IN model for different parametrizations *θ*_*i*_ with stimuli *I*_Stim_ (normalized in gray). Solutions for *v*(*t*) and the respective spike-times for a reference solver (black) and the original solver scheme (orange). For plotting, *v*(*t*) were clipped at 30. For both solutions, spike-times are shows in a raster plot (bottom) together with spike-times of 20 samples from a 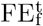 solver (blue).

**Figure S2:**
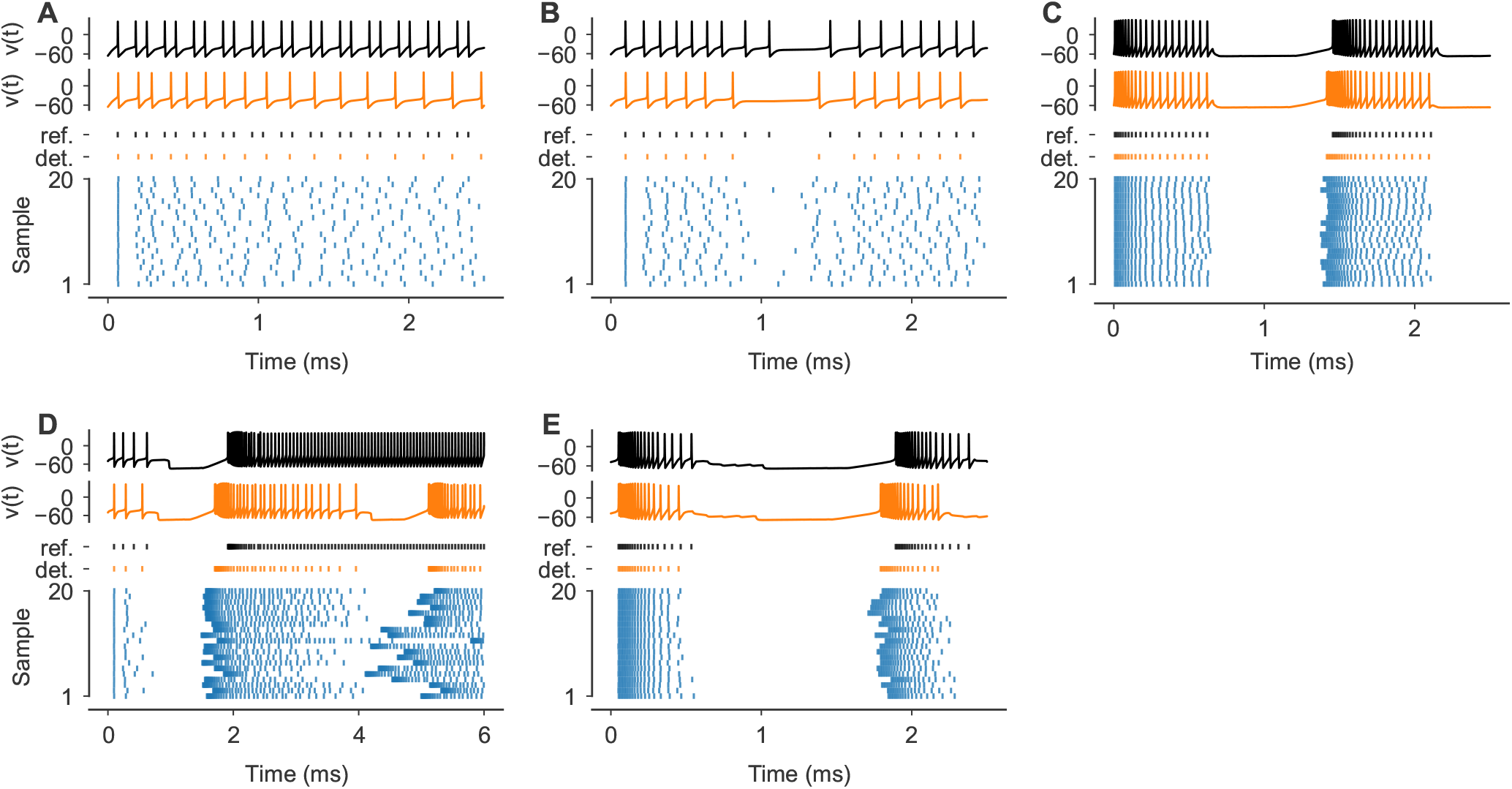
**(A-E)** As in Figs. 2D and 2E. Simulations of the STG model for all five synaptic parametrizations (see Appendix C) from 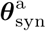 to 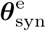, respectively. Solutions for the membrane potential *v*(*t*) of the LP neuron are shown for a reference solver (black) and a deterministic EE solver with Δ*t*=0.1 ms (orange). *Bottom panels*: Spike-times of the LP neuron are shown for the reference (black), the deterministic EE solution (orange) and for 20 samples from a probabilistic 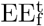 with Δ*t*=0.1 ms (blue).

**Figure S3:**
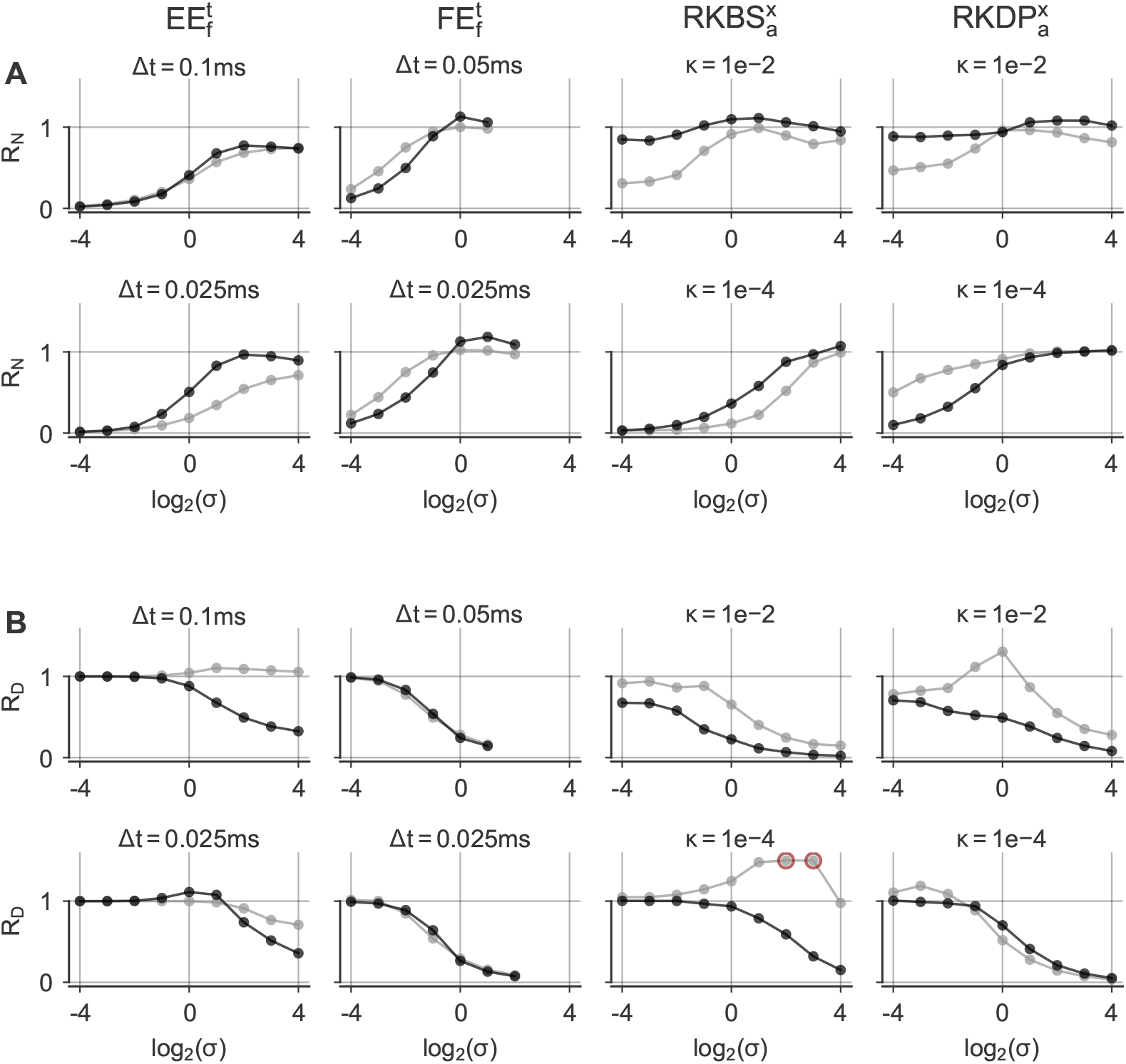
**(A, B)** MAE ratios *R*_*N*_ and *R*_*D*_ for different solvers (column titles) and step-sizes Δ*t* / tolerances *κ* (panel titles) for the step (grey) and the noisy step (black) stimulus as a function of the perturbation parameter *σ* (x-axis), respectively. See also Fig. 6D. Values larger than 1.5 were clipped (indicated by red circles).

